# Interplay of Proactive and Reactive Control in Language Production

**DOI:** 10.64898/2026.07.09.737628

**Authors:** Katherine D. Andrade, Dakota L. Melton, Stephanie K. Riès

## Abstract

Language production requires the coordination of multiple cognitive processes. The ability to anticipate and override a habitual response in favor of a contextually-appropriate response are key subprocesses of cognitive control which enable speakers to communicate effectively. Word retrieval involves the co-activation of semantically related alternatives from which the speaker must select the appropriate target representation. Although cognitive control mechanisms have been proposed to contribute to resolving semantic interference during language production, the nature of these control processes remain unclear. Studies investigating the temporal dynamics of cognitive control during decision making tasks have led to a distinction between two operating processes: proactive control, initiated prior to the occurrence of conflict, and reactive control recruited after conflict is detected. We investigated the roles of proactive and reactive control in resolving interference between competing linguistic representations during word retrieval. We analyzed congruency sequence effects combined with delta-plot distributional analyses to dissociate potential adjustments in proactive versus reactive cognitive control in a picture-naming task manipulating semantic context compared to a minimally-linguistic Stroop-like paradigm. Reaction time distributional properties following semantically related trials revealed the engagement of proactive control in semantic interference resolution during word retrieval in the PWI task. In contrast, reactive inhibitory control was engaged in resolving semantic interference following low conflict trials. This distinction was not present in the minimally-linguistic task, which did not appear to engage adaptive control to the same extent. These findings demonstrate that both proactive and reactive cognitive control mechanisms contribute to language production, and are engaged dynamically, adjusting trial-by-trial to resolve semantic interference during word retrieval. In addition, our study provides important insight into the comparison of language with other cognitive domains and positions linguistic paradigms as being instrumental in the study of cognitive control dynamics.

## 1. INTRODUCTION

For most individuals, language comes naturally and occurs seamlessly. Despite the ease with which we communicate, language production has been proposed to rely on highly efficient cognitive control mechanisms (Ferreira & Pashler, 2002). Cognitive control encompasses multidimensional operations, including goal-maintenance, response inhibition, selective attention, conflict resolution, task switching, and error monitoring, required to optimize performance (Miller & Cohen, 2001, Miyake et al., 2000). These processes have been extensively studied in other cognitive domains (i.e., memory, Engle, 2018, motor control, Georgopoulos, 2002, and executive control, Diamond, 2013). However, the multidimensional aspects of cognitive control remain relatively understudied in language production. When planning to produce a word or multi-word phrase, speakers must maintain the goals of the conversation in mind while simultaneously integrating new information into working memory to guide an appropriate oral response (Acheson & MacDonald, 2009). The spread of activation within a densely interconnected system simultaneously activates multiple lexical representations. Control mechanisms have been proposed to be required to resolve potential conflict between the target item and co-activated representations in order to appropriately select and produce the intended response (Freund et al., 2016). This study sheds light on the nature of cognitive control processes involved in resolving this conflict.

### 1.1. Non-linguistic cognitive control

#### 1.1.1. Within-Trial Conflict Resolution

A variety of conflict tasks have been designed to investigate cognitive control, including the Eriksen Flanker task (Eriksen & Eriksen, 1974), the arrow version of the Flanker task (Stoffels & van der Molen, 1988), the Simon task (Simon & Rudell, 1967), the Stroop task (Stroop, 1935), or the spatial Stroop task (Liu et al., 2004, Giezen et al., 2015), to name a few. In these tasks, participants are required to respond quickly typically by pressing left or right buttons depending on a characteristic of the stimulus. For example, the spatial Stroop task requires participants to press left or right buttons depending on the direction of an arrow on the screen while ignoring the position of the stimulus relative to a central fixation cross. During congruent trials, the irrelevant dimension of the stimulus (i.e., its position relative to the fixation cross) matches the relevant dimension of the stimulus (i.e., the direction the arrow is pointing to), and the correct response is therefore easily chosen. In incongruent trials however, the irrelevant dimension of the stimulus does not match the relevant dimension of the stimulus and the correct response is therefore more difficult to choose. The fact that responses in incongruent trials are slower and more error prone compared to congruent trials is referred to as the congruency effect (e.g., Ericksen & Ericksen, 1994, Stroop, 1935, Simon & Rudell, 1967). The origin and magnitude of the congruency effect can vary depending on the task performed, as presented in Kornblum’s dimensional overlap model (Kornblum et al., 1990), but cognitive control is typically needed to resolve the conflict arising from these congruency manipulations.

#### 1.1.2. Between-Trial Conflict Resolution

In addition to playing a key role in resolving conflict within a trial, cognitive control can be adjusted between trials. Indeed, performance on a current-trial has been shown to be influenced by the presence of conflict in the previous-trial. This phenomenon is referred to by various names-such as the congruency sequence effect (CSE), the Gratton effect, conflict adaptation effect, or the sequential modulation effect (Gratton et al., 1992, Braver, 2012, Egner, 2007, Roquet et al., 2020). Specifically, when a congruent trial is preceded by an incongruent trial (iC), performance tends to be lower compared to when a congruent trial is preceded by another congruent trial (cC). Conversely, an incongruent trial that is preceded by another incongruent (iI) trial typically leads to better performance compared to when an incongruent trial is preceded by a congruent trial (cI). One of the main models used to explain the CSE is the conflict-monitoring account (Botvinick et al., 2001), according to which an internal monitoring system detects conflict and determines the need for increased control to facilitate interference resolution in the next trial (but see Egner, 2014). Freund and Nozari (2018) build upon the conflict-monitoring account by suggesting that the system can increase control to bias processing towards the target dimension or suppress processing of the distractor dimension to improve performance on the following trial when response conflict occurs. Therefore, the CSE has been used to provide a measurable signature of how the cognitive system adapts to conflict on a trial-by-trial basis (Braem et al., 2019) and can be used to assess modes of cognitive control that differ in temporal dynamics.

#### 1.1.3. The Dual Mechanisms of Cognitive Control

The dual mechanisms of cognitive control model creates a distinction between two operating inhibitory control processes: proactive and reactive control (Braver, 2012, Meyer & Bucci, 2016). According to Braver (2012), proactive control is initiated before conflict arises and can be viewed as a form of attention enhancement by which information relevant to the task goal is actively maintained to bias attention, perception, and action systems in an objective-driven approach (Miller & Cohen, 2001, Braver et al., 2007). In contrast, during reactive control, attention is recruited after conflict is detected and can be viewed as a ‘late correction’ mechanism (Braver et al., 2007). Researchers have designed paradigms to manipulate proactive and reactive biasing at the local and global level to differentiate their temporal dynamics (Kricheldorff et al., 2023, Xiang et al., 2016). The CSE has been argued to be usable as a local measure of proactive control, particularly in trial sequences in which adjustments in control are made directly following conflict from the previous trial in preparation for the current response (Egner, 2007, Kricheldorff et al., 2023). Instead, reactive control reflects transient adjustments capturing just-experienced conflict between competing response representations (Braem et al., 2019, Kricheldorff et al., 2023) and can be measured when there is no possibility for adjustment in control from one trial to the next. Although, some studies have argued that the CSE is a result of the upregulation of reactive control from the previous trial (Duthoo et al., 2014, Duthoo & Notebaert, 2012, Egner et al., 2010, Scherbaum et al., 2012, Yang & Pourtois, 2022).

Distributional properties can reveal information beyond average reaction time (RT) and accuracy measures (Luce, 1991). Specifically, distributional analysis can uncover how an experimental effect, such as the congruency effect, changes throughout the distribution of reaction times (Balota & Yap, 2011) using Vincentile (Vincent, 1912) or delta plots (De Jong et al., 1994). These analyses have revealed that the size of congruency effects can increase or decrease with increasing RTs depending on the selected cognitive paradigm (Tang et al., 2022). In many tasks eliciting conflict, the congruency effect increases with increasing reaction times. The interpretation of such an increasing effect is due to the prepotent task-irrelevant attribute becoming more active with time, making the decision process take longer to happen (Miller and Schwarz, 2021). The activation-suppression model (Ridderinkhof, 2002) explains the congruency effect as arising from differences in top-down inhibition of the task-irrelevant response. Specifically, inhibitory control builds up over time and can attenuate congruency effects in trials with slower responses more so compared to faster responses (Ridderinkhof et al., 2004, but see Schwarz & Miller, 2012). When inhibition is strong to begin with, as is the case in a system where conflict is expected to happen, the congruency effect can be reduced earlier in the RT distribution (Ridderinkhof et al., 2002). This is the pattern that we would expect in a system that relies more on proactive than reactive control. However, when inhibition is weak to begin with, congruency effects grow with increasing RTs initially, but can be reduced later in the RT distribution once inhibition has accrued (Ridderinkhoff et al., 2002; Xiang et al., 2016). This is the pattern we would expect in a system that relies primarily on reactive control. As such, both control mechanisms may operate in tandem to reduce interference at different timescales, with proactive control acting earlier and reactive control emerging later in the RT distribution. Distributional analyses can therefore contribute to the understanding of the temporal dynamics of cognitive control involved in conflict adaptation (i.e., top-down increase of control following conflict detection) and conflict resolution during response selection.

### 1.2. Cognitive control in language production

#### 1.2.1. Resolving Conflict Between Semantically-Related Representations

The ability to anticipate and resolve conflict are key subprocesses of cognitive control that enable speakers to communicate efficiently. In general, it has been suggested that cognitive control plays a key role in language production to resolve interference from co-activated representations during word retrieval, between semantically-related representations. Researchers have used a variety of paradigms to study semantic interference, including: the picture-word interference (PWI) paradigm (Lupker, 1979, Schriefers et al., 1990), the blocked cyclic naming task (Belke et al., 2005), the continuous naming paradigm (Howard et al., 2006), or semantic task-switching paradigms (Viviani et al., 2025). In the PWI task, participants respond by verbally naming a target picture, while ignoring a superimposed distractor word. When the picture name and the distractor word are semantically-related in a taxonomic way (e.g., pear and apple), a semantic interference effect will typically be observed on performance: Reaction times will be longer and accuracy will be lower in trials in which the picture name and distractor word are semantically-related compared to trials in which the picture name and distractor word are unrelated. One of the most influential models explaining the semantic interference effect during word retrieval is Oppenheim et al.’s (2010) incremental learning model. According to this model, word retrieval is achieved by strengthening the connections between the semantic and lexical features of the target word, while weakening the connections between the semantic and lexical features of competitors. When the activation of representations that are semantically-related to the target is increased, a “booster” mechanism may be engaged to facilitate the selection of the target representation if there is sufficient differentiation between co-activated representations. This “booster” mechanism has been proposed to reflect cognitive control involved in resolving the conflict induced by semantically-related items during word retrieval and to be analogous to the cognitive control and monitoring model proposed by Botvinick et al., (2001). According to this model, the monitoring mechanism tracks the degree of activation levels between representations, and selects an item once its activation surpasses a certain threshold. Cognitive control is implemented when conflict between possible responses makes reaching the selection threshold difficult. Nozari & Hepner, (2019) proposed that this selection threshold is flexibly modulated and a more recent proposal by (Nozari, 2025) suggests that the need for cognitive control may be limited to specific cases where a prepotent response needs to be inhibited. This would be the case in the PWI and spatial Stroop task above-mentioned.

#### 1.2.2. Mechanisms of control in language production

Trial-by-trial adjustments also exist in language production (Duthoo et al.,2014, Freund et al., 2016, Shitova et al., 2017, demonstrating that single-word production can be monitored and adjusted from one trial to the next, similar to within non-linguistic tasks (Ericksen & Ericksen, 1994, Simon & Rudell, 1967). Shitova et al., (2017) investigated the neural bases of the CSE in a PWI task by measuring ERPs and oscillatory power. They reported the expected quantitative differences on error rates and RTs following semantically related trials in addition to an N400 effect and frontal theta power modulations consistent with the recruitment cognitive control, though its nature remains to be specified (see also Piai & Zheng, 2019). Semantically-related trials can be seen as analogous to incongruent trials, while identical trials (i.e., when the distractor word matches the picture name) can be seen as analogous to the congruent condition in non-linguistic tasks. Therefore, it is not surprising that adjustments in control can be made from one trial to the next in language production similarly as in non-linguistic control tasks.

Similarly, as in non-linguistic cognitive control studies, the distributional characteristics of the semantic interference effect on RTs have been studied and are argued to inform underlying cognitive control processes (Shao et al., 2013). Studies using the PWI paradigm have quantified individual and group inhibitory control dynamics by correlating the magnitude of the semantic interference effect to the slope of the delta plot for the slowest RTs (Shao et al., 2013, Shao et al., 2015, Roelofs et al., 2011). Shao et al., (2013) reported that individuals with more negative slopes in the slowest quantiles of the delta plot exhibited smaller semantic interference effects, suggesting that these individuals’ ability to resolve semantic interference was linked to their greater efficiency in the application of selective inhibition. However, the differential involvement of proactive versus reactive control in language production has received far less attention than outside of language. Thus, distributional changes in the time course of the semantic interference effect require further investigation and may provide insight into the specific modes of cognitive control processes engaged to resolve semantic interference.

### 1.3. Current Study

In the present study, we investigate how proactive and reactive control mechanisms contribute to the resolution of semantic interference in language production. By investigating trial-by-trial performance adjustments and distributional characteristics of reaction times, this study aims to identify which cognitive control mechanisms are implemented to adapt to and resolve conflict within and outside of language. To this end, we examine the semantic interference and Stroop effects (i.e., congruency effects) in healthy adult participants to reveal proactive and reactive control dynamics across two tasks: a linguistic picture-word interference and a minimally-linguistic spatial Stroop task. The PWI and spatial Stroop tasks (Liu et al., 2004, Giezen, 2015, Blumenfeld & Marian, 2014) have been used concurrently in studies assessing bilingual language control and non-linguistic inhibitory control (Andrade et al., 2025, Gálvez-McDonough et al., 2025). Several of these studies have found parallels between linguistic and non-linguistic interference effects. However, whether or not proactive and reactive control are engaged similarly to resolve semantic interference when we speak compared to non-linguistic decision making is unknown. We hypothesize that increase demands in reactive control will be associated with larger Stroop effects with increasing response times. Meanwhile, the engagement of proactive control will lead to smaller Stroop effects on trials following incongruent trials as the detection of conflict at trial n-1 attenuates the cost of processing subsequent conflict. Stroop effects will be maximal for the shorter response times and progressively decrease throughout the distribution, with the greatest reduction at longer RTs as shown in negatively sloped delta plots (Wylie et al., 2010, Xiang et al., 2016). We hypothesize that these effects will be comparable across the PWI and spatial Stroop task and correlated in magnitude. By investigating how proactive and reactive control unfold while we speak, we aim to shed light on the complex cognitive processes supporting how individuals maintain fluency and coherence in communication despite the potential for semantic interference.

## 2. METHODS

### 2.1. Participants

Thirty-six (25 females, mean age: 30.78 years, SD: 11.14, mean education:17 years, SD: 1.7) native English speakers between the ages of 18-60 years old^i^ participated in the study. This age-range was chosen as this group constitutes an age-matched control group for another study. All participants had no history of neurological impairment, hearing loss, and had normal or corrected-to-normal vision. The data from 4 participants were excluded from analysis due to technical issues during data collection. Therefore, analysis was performed on the remaining 32 participants (22 females, mean age: 30.44, SD:10.75, mean education: 15 years, SD: 1.7). All participants provided written consent and completed neuropsychological assessment to test their cognitive function, linguistic knowledge and reading skills (see supplementary materials section S1 and table S1).

### 2.2. Materials

Participants performed a picture-word interference task (PWI, e.g., Lupker, 1979) and an arrow version of the Stroop task (i.e., spatial Stroop task; Giezen et al., 2015). The PWI task is an established task used to assess lexical access (Dell’Acqua et al., 2007, Shao et al., 2015). The semantic version of the task elicits the semantic interference effect: where performance is lower if the picture name and distractor word are from the same taxonomic semantic category (e.g. cat and dog are both animals) compared to when they are semantically unrelated. The spatial Stroop task is an alternative version of the Stroop task (Liu et al., 2004; Giezen, 2015; Blumenfeld and Marian, 2014). Participants are required to respond to the direction of a left-pointing or right-pointing arrow, while ignoring its spatial location. Incongruency between the arrow’s direction and spatial location, such as a right-pointing arrow presented on the left side of a central fixation cross, results in slower RTs and decreased accuracy compared to congruent trials, where the direction and side of presentation of the arrow are matching (MacLeod, 1991, Giezen, 2015). Responding to the direction of an arrow while ignoring its spatial location in the spatial Stroop task or naming the color of a word while ignoring what the word is, requires cognitive control to resolve the interference between task-relevant and task-irrelevant features. The instruction to attend to the arrow’s direction or the picture’s name is activated in a controlled manner and can conflict with the more automatic processing of responding to the spatial location or the reading of the distractor word, i.e. the prepotent response (Matsushima (Matsushima & Aznar-Casanova, 2025)et al., 2025). The decision to be made in this spatial Stroop task requires little to no lexical access (i.e. only two possible responses – “Right” or “Left” dependent on the arrow direction) in comparison to naming pictures in the PWI task which requires choosing between several possible activated lemmas. Previous studies have used these two tasks to investigate potentially overlapping control processes in linguistic and non-linguistic domains (Gálvez-McDonough et al., 2024; Andrade et al., 2025). Here, we will use verbal rather than manual responses to maximize compatibility across paradigms, and to ensure that any differences between tasks are not attributable to response modality.

### 2.3. Design and Stimuli

The current study followed a 3 x 2 design, with trial type and task as within-subject factors. For the PWI task, participants were tasked to name pictures while ignoring the superimposed distractor word belonging to three conditions: semantically related, semantically unrelated, and identity. The stimuli consisted of 60 colored photographs with above 80% naming agreement issued from the BOSS database (Brodeur et al., 2014). Each target stimuli was presented 3 times, once per condition (i.e., incongruent, congruent, and neutral), throughout the entire list. Three lists of 180 target-distractor word pairs were developed to avoid systemic order effects. The order of stimulus presentation was pseudorandomized to ensure that: (1) targets belonging to the same semantic category were separated by stimuli from at least two other categories; (2) identical pictures were separated by at least three other stimuli; (3) distractor words were separated by at least three stimuli containing different distractor words; (4) the stimulus condition did not repeat more than three times consecutively; and (5) the initial phonemes of picture names and distractor words were controlled for, such that neighboring stimuli did not share the same initial phonemes. Embedded within the traditional PWI task, we controlled for the sequential order of trial types, such that all possible previous-current trial congruency combinations occurred a similar number of times throughout the experiment. From here on, the semantically-related trials will be referred to as *incongruent,* the identity trials as *congruent* to align with the spatial Stroop task. Semantically-unrelated trials will continue to be referred to as *unrelated* and will correspond to the *neutral* trials in the spatial Stroop task.

The spatial Stroop task was administered to examine participant’s cognitive control abilities in a minimally linguistic language production task. Participants were tasked to name the direction the arrow was pointing to while ignoring its spatial location. Stimuli consisted of white colored left-or right-pointing arrows displayed on a black background and in 3 conditions: *congruent*, *incongruent*, *neutral* (see Giezen, 2015). There was an equal proportion of left-and right-pointing arrows in each condition. Stimuli were presented across 6 blocks, with 30 trials per block, adding up to a total of 180 trials. Three lists were developed to account for systemic order effects. To examine sequential adjustments in cognitive control, the sequence of trial types was pseudorandomized such that all possible previous-current trial congruency combinations occurred throughout the experiment as in the PWI task (i.e., cC, cI, cU, iC, iI, iU, uC, uI, uU).

### 2.4. Procedure

The study was conducted in a single hour and a half session. After giving consent and completing demographic background and neuropsychological assessment, individuals participated in the two verbal behavioral tasks. The experiment was run on Presentation software (version 18.0, Neurobehavioral Systems, Inc., Berkeley, CA). Stimuli were presented at the center of a 16 in. Dell monitor in front of participants. Verbal responses were recorded using a Marantz professional MPM-1000 condenser microphone and manually transcribed offline using CheckVocal (Protopapas, 2007), which was also used to mark vocal onset.

The order of task administration was counterbalanced across participants. Prior to the start of the PWI task, participants were familiarized with all the pictures used in the experiment by reviewing a slideshow of each picture with its expected picture name written underneath. At the start of each task, participants completed a practice block that was excluded from analysis. Participants were asked to name the pictures or name the direction of the arrow as fast and accurately as possible. For both tasks, response times for each trial reflect the difference between the onset of stimulus presentation and the time of vocal onset, with a timeout value set to 3000ms. In the PWI task, each trial consisted of the following: (1) a 300-750ms jittered fixation cross (2) the stimulus for 2000ms, and (3) a white blank screen for 2000ms. In the spatial Stroop task, each trial consisted of the following: (1) a 300-750ms jittered fixation cross; (2) the stimulus for 700ms; (3) a black blank screen for 2000ms. Breaks were self-paced between trial blocks.

### 2.5. Analysis

Accuracy and reaction times were analyzed using R (R Core Team, 2020) and Rstudio (Rstudio Team, 2020), including the lme4 (Douglas et al., 2015), lmerTest (Kuznetsova et al., 2017), car (Fox and Weisberg, 2019), dplyr (Wickham et al., 2023) packages. The first trial of every block was removed as there are no data points from the preceding trial. We analyzed the accuracy data using logistic mixed-effects models (Baayen et al., 2008, Jaeger, 2008). We tested for the main effects of current trial congruency, previous trial congruency, task, and their interaction as within-participant factors. The model included random intercepts for participant and item, as well as by-participant random slopes for current-trial congruency, previous-trial congruency, and their interaction. For the analysis of RTs, only correct trials were analyzed. All incorrect trials (PWI: 5.39%, spatial Stroop: 1.17% of all trials), omissions (PWI: 0.86%, spatial Stroop: 0.18% of all trials), and RT outliers (±3 SDs within subject; PWI: < 4% and spatial Stroop: <15% of correct trials) were excluded. Correct trials were defined as responses matching the stimulus name but semantically identical names for an item were accepted as correct (e.g., bike-bicycle, bunny-rabbit, etc.). Vocal responses that included anything besides the one-word expected response was considered as incorrect (e.g., incomplete utterances, stuttering, hesitations, etc.). RTs were log-transformed to reduce skewness and approach a normal distribution. Within each of the two tested paradigms, a four-step analysis was computed: (1) linear mixed-effect model analyses on RTs, (2) Vincentile distributional analysis on bin-wise mean RTs, (3) slope analysis on delta plot distributions, and (4) correlational analysis to compare the effects of interest in both tasks. Analyses were performed on subsets of the data to compare congruent trials to incongruent trials (i.e., Stroop effects, part 1) and to compare incongruent trials to unrelated/neutral trials (i.e., Interference effects, part 2) in and across the two paradigms.

Distributional analyses were conducted using the Vincentile method (Vincent, 1912, Ratcliff,1979). RTs were first rank-ordered from fastest to slowest separately for each CSE condition (i.e. cC, cI, cU, iC, iI, iU, uC, uI, uU) within each participant (18-22 trials each). RTs were then divided into five quintiles (as in Shao et al., 2013, Shao et al., 2015, Fuhrmeister & Bürki, 2022). Quintile-based distributional methods, such as Vincentizing, is effective when there are as few as 10-20 RTs per participant per condition (Ratcliff, 1979, Jiang et al., 2004), while parametric fitting requires a larger number of trials within each condition for distributional estimates. Therefore, Vincentile analyses were applied to create group RT distributions per condition per task. For each task, we analyzed the effects of current trial congruency, previous trial congruency, bin, and their interaction using repeated-measures by-participants ANOVAs.

Delta plots (De Jong et al., 1994) were then created to estimate the time course of conflict adaptation. For each participant, we calculated the difference (delta Δ) between conditions within each bin (i.e. Stroop Effect: ΔIIvIC: iI – iC, ΔCIvCC: cI – cC; Interference effect: ΔIIvIU: iI – iU, ΔUIvUU uI-uU). For each participant, linear slopes were estimated by regressing each delta measure on bins, yielding participant specific slope estimates across the RT distribution. We then tested whether slopes differed by condition. The effect of condition on slopes was analyzed using repeated-measures by-participant ANOVAs. Additionally, considering inhibitory control has been shown to accumulate over time and affect longer responses the most (see introduction), a post-hoc paired student t-test was conducted to compare the slope of the inflection between the last delta segment (bin 4-5) and the slope of the earlier delta segments (bins 1-4). Finally, participant-specific Stroop and interference effect coefficients were extracted from the linear mixed-effect models for each task. Pearson correlations were conducted to assess the correlation between Stroop effects and interference effects between the PWI task and the spatial Stroop task. Bonferroni corrections were applied to control for family-wise errors.

## 3. RESULTS

### 3.1. Part 1: Stroop Effect

#### 3.1.1. Stroop and Congruency Sequence Effect on Accuracy

In both the PWI task and the spatial Stroop task, participants demonstrated high accuracy (Table 2). There was a significant main effect of current trial congruency, χ²(1)= 8.8, p=.003, and a main effect of task, χ²(1)= 4.86, p=.027, on accuracy. Participants were more accurate when the current trial was congruent compared to incongruent (β_raw_=-0.62, SE= 0.21, z=-2.97, p= 0.003) and more accurate in the spatial Stroop task compared to the PWI task (β_raw_=-0.60, SE= 0.27, z=-2.21, p= 0.027). There was no significant main effect of previous trial congruency, χ²(1)= 2.96, p= 0.085. Additionally, there was no significant interaction between previous and current trial congruency, χ²(1) = 3.25, p=.071, previous trial congruency and task, χ²(1)= 0.33, p=.563, current trial congruency and task, χ²(1,64)= 0.10, p=.756, nor a three way interaction between previous trial congruency, current trial congruency, and task, χ²(1)= 0.01, p= 0.925, on accuracy. This indicated that there was no congruency sequence effect on accuracy and this did not differ between tasks, potentially because accuracy was so high overall.

**Table 2:**
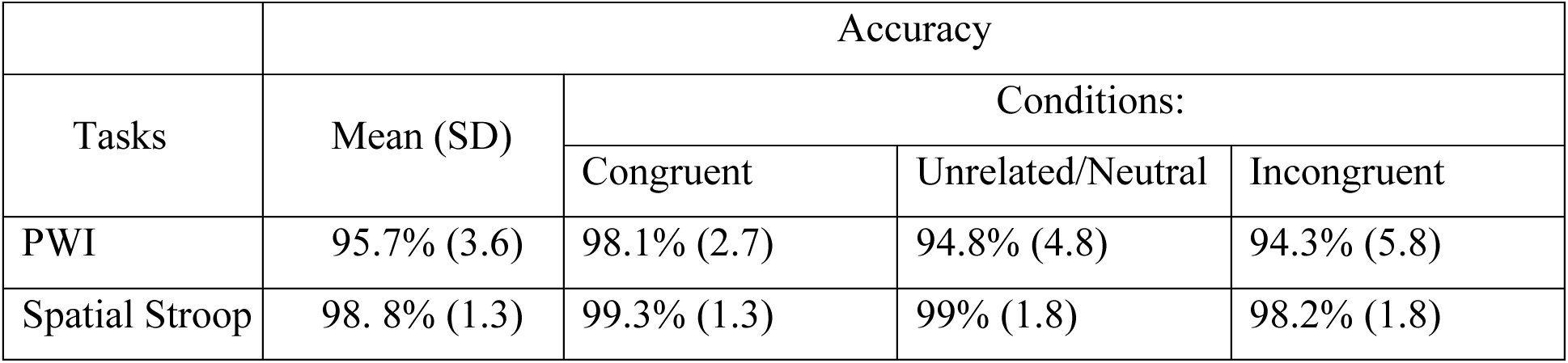
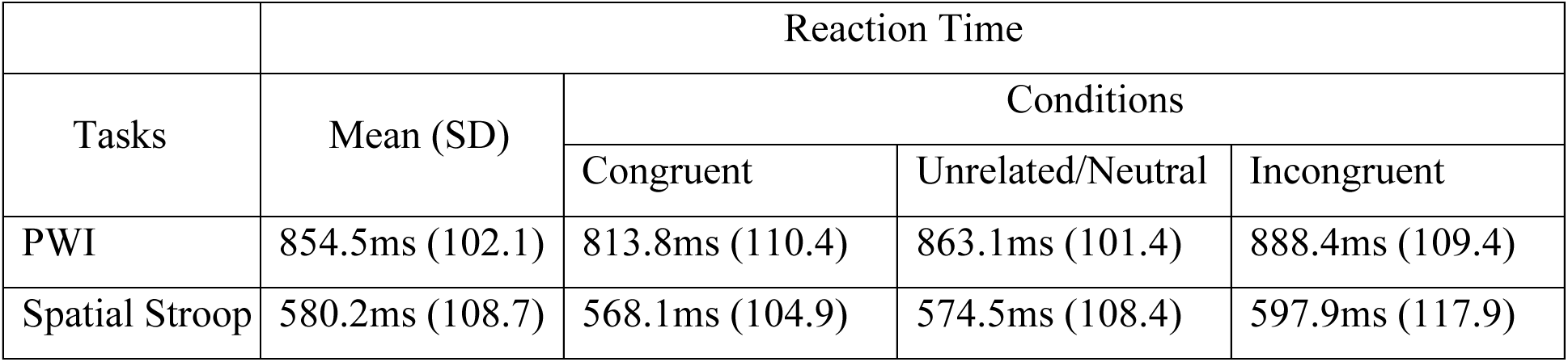
Mean accuracy rates and reaction time per task and condition.

#### 3.1.2. Stroop and Congruency Sequence Effect on Reaction Times

There was a significant main effect of current trial congruency, χ²(1)= 91.60, p< 0.001, and a main effect of task, χ²(1)= 77.9, p< 0.001. Participants were significantly slower to name pictures in the incongruent condition compared to congruent condition (β_raw_= 5.59×10^-2^, SE= 5.83×10^-3^, t= 9.57, p< 0.001) and performed more slowly in the PWI task compared to the spatial Stroop task (β_raw_=-3.70×10^-1^, SE= 4.19×10^-2^, t= 8.83, p< 0.001). There was a significant interaction between previous trial congruency and current trial congruency, χ²(1) =16.85, p<.001, indicating that the current trial Stroop effect was reduced following an incongruent trial. Additionally, there was a significant interaction between current trial congruency and task, χ²(1)= 29.3, p< 0.001, indicating that the current trial Stroop effect was greater in the PWI task compared to the spatial Stroop task (β_raw_=-4.09×10^-2^, SE= 7.56×10^-3^, t=-5.41, p< 0.001). There was no significant main effect of previous trial congruency, χ²(1)= 0.69, p= 0.41. There was also no significant interaction between previous trial congruency and task, χ²(1)= 2.73, p= 0.099, nor was there three-way interaction between previous trial congruency, current trial congruency, and task, χ²(1)= 1.93, p= 0.165, indicating that the magnitude of the modulation of the Stroop effect by previous trial congruency did not differ between the PWI and the spatial Stroop task. (β_raw_= 8.02×10^-3^, SE= 5.78×10^-3^, t= 1.39, p= 0.165).

#### 3.1.3. PWI Task-Vincentile Analysis

A repeated-measures ANOVA revealed a significant main effect of current trial congruency on mean-binned RT, *F*(1, 31) = 44.17, *p*< 0.001, *η*p²= 0.59. As expected, there was a significant main effect of bin, *F*(1, 31) = 615.42, *p*< 0.001, *η*p²= 0.95. Additionally, there was a significant three-way interaction between previous trial congruency, current trial congruency, and bin (*F*(1, 31)= 4.79, *p*= 0.036, *η*p²= 0.137), indicating that the CSE increased with increasing RTs (as shown in Figure 2). There was no significant main effect of previous trial congruency, *F*(1, 31)= 1.93, *p*= 0.175, *η*p²= 0.06, and no significant interaction between current trial congruency and previous trial congruency, *F*(1,31)=2.68, *p=* 0.112, current trial congruency and bin, *F*(1, 31)= 2.68, *p*= 0.112, *η*p²= 0.08, or previous trial congruency and bin, *F*(1, 31)= 0.032, *p*= 0.86, *η*p²< 0.001).

**Figure 1.**
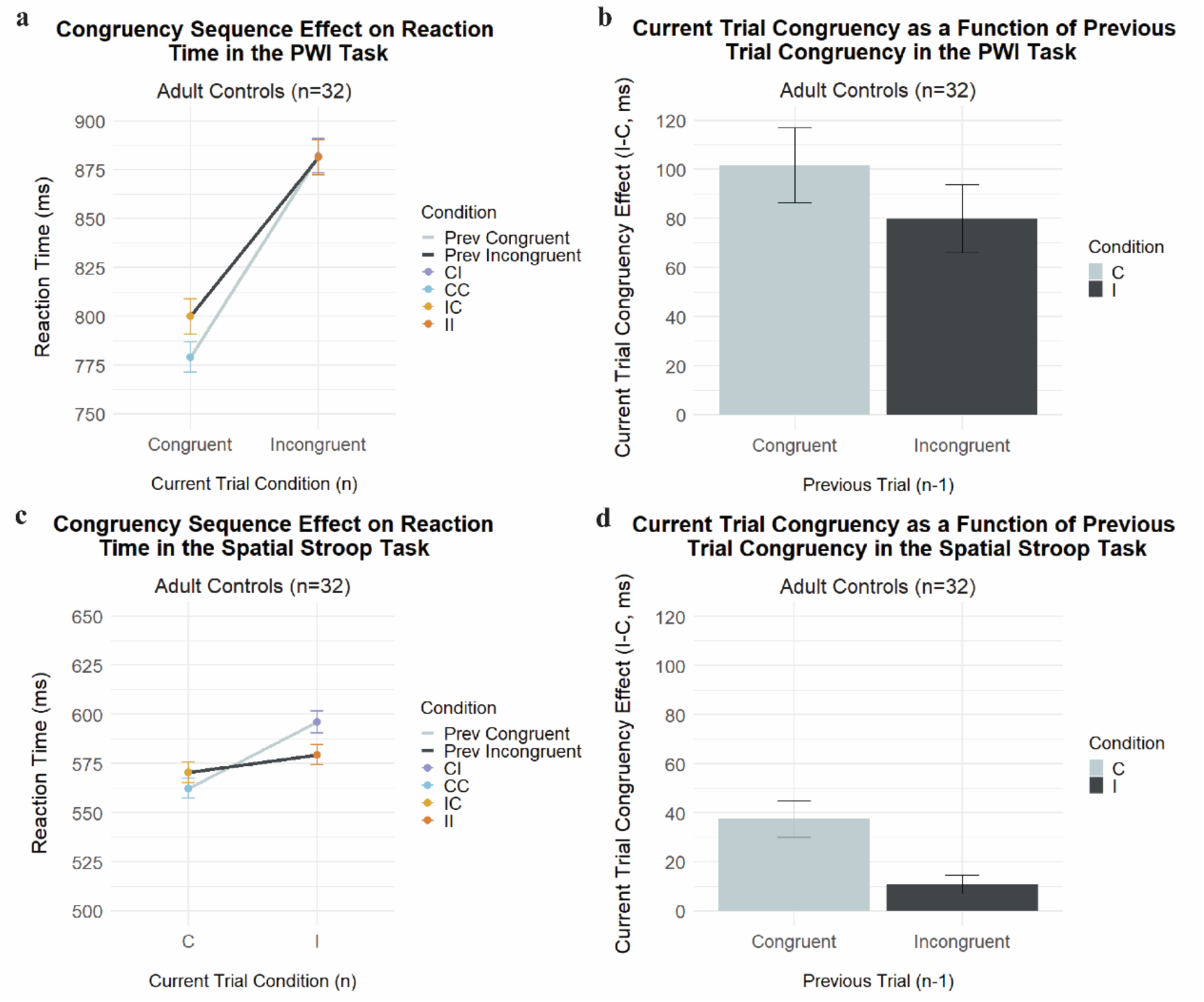
*Left-* CSE on reaction times in adult controls in the PWI (a) and spatial Stroop task (c) in four conditions (i.e., cC-blue, cI-purple, iC-yellow, iI-orange). Previous congruent trials are the cooler colors (blue and purple) and are connected by a light grey solid line. Previous incongruent trials are the warmer colors (yellow and orange) and are connected by a dark grey solid line. *Right-* Difference between mean RTs in incongruent versus congruent trials as a function of previous trial condition (congruent, incongruent), illustrating a reduced Stroop effect following incongruent trials relative to congruent trials in the PWI (b) and spatial Stroop (d) task.

**Figure 2.**
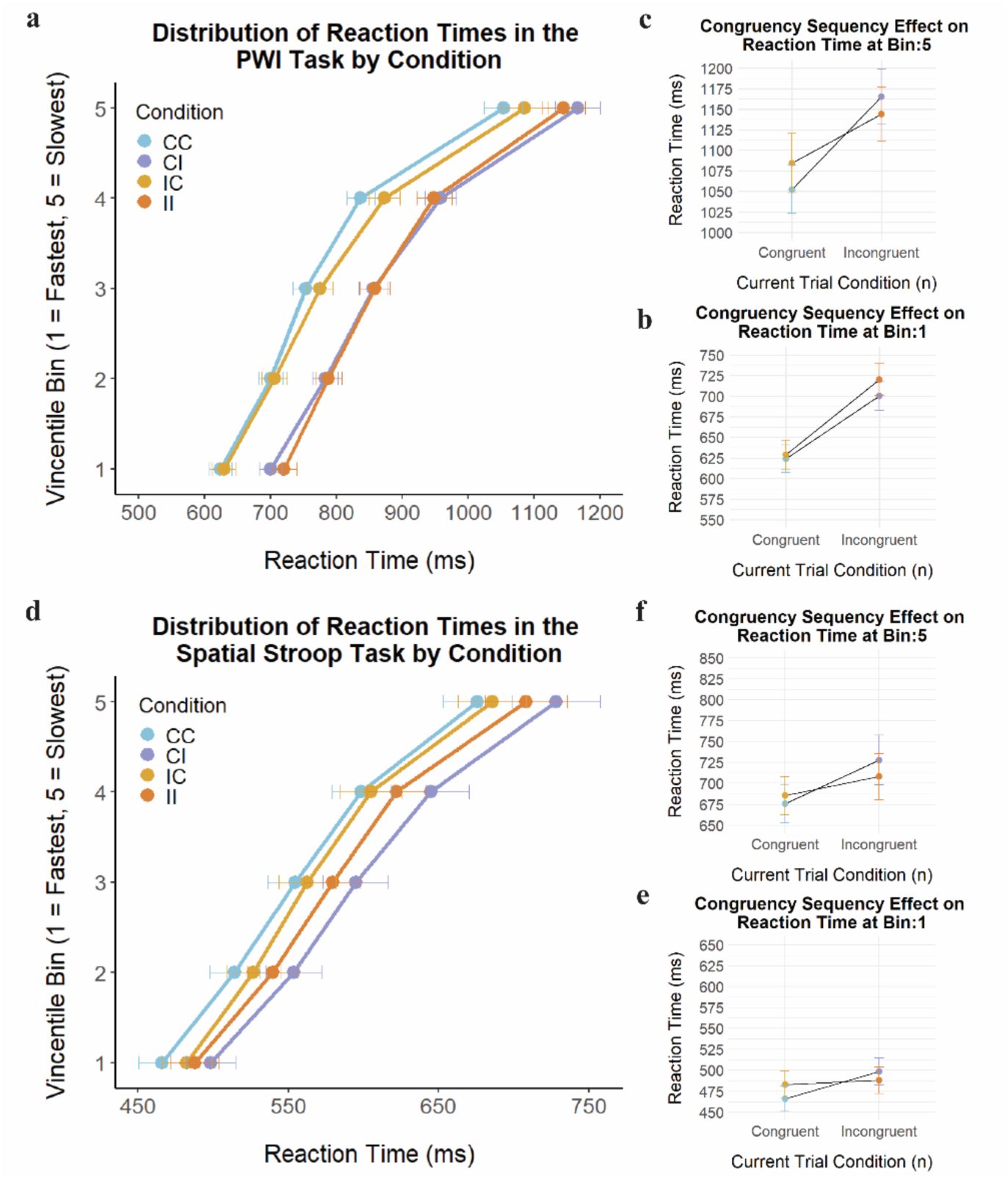
*Left-* Vincentile plots demonstrating the distribution of reaction times across quintile (1-fastest, 5-slowest) by condition (i.e., cC-blue, cI-purple, iC-yellow, iI-orange) in the PWI task (a) and spatial Stroop task (d). Error bars denote the standard error of the mean by condition, and bin. *Right-*CSE on reaction time in the PWI (top) and spatial Stroop task (bottom), these plots highlight the evolution of the CSE across the RT distribution from bin 1 (b, e) to bin 5 (c, f).

#### 3.1.4. Spatial Stroop Task-Vincentile Analysis

A repeated-measures ANOVA revealed a significant main effect of current trial congruency, *F*(1, 31)= 43.79, *p*< 0.001, *η*p²= 0.59. As expected, there was a significant main effect of bin, *F*(1, 31) = 287, *p* < 0.001, *η*p²= 0.90. Additionally, there was a significant interaction between current trial congruency and previous trial congruency, *F*(1,31)= 11.08, *p*= 0.002, *η*p²= 0.26. There was no significant main effect of previous trial congruency, *F*(1, 31)=.84, *p*= 0.368, *η*p²= 0.03, no significant interaction between previous trial congruency and bin, *F*(1, 31)= 1.70, *p*= 0.20, *η*p²= 0.05, current trial congruency and bin (*F*(1,31)= 3.22, *p*= 0.083, *η*p²= 0.09), nor a three-way interaction between previous trial congruency, current trial congruency, and bin, *F*(1, 31)= 0.037, *p*= 0.849, *η*p²< 0.001, indicating that the magnitude of the CSE did not change across the RT distribution in the spatial Stroop task.

#### 3.1.5. PWI Task-Delta Slope Analysis

For each participant, we calculated the difference (delta, Δ) between conditions within each bin (i.e. Stroop Effect: ΔIIvIC: iI – iC, ΔCIvCC: cI – cC). We examined the slope of these delta plots which capture the evolution of the difference between conditions across the RT distribution. A repeated-measures ANOVA revealed a significant main effect of Condition on the delta plot slope*, F*(1, 31) = 4.79, *p* = 0.036, *η*p² = 0.13. The slope of the delta plot for iI vs. iC conditions (i.e., ΔIIvIC) was negative (mean=-6.99 ms), indicating that the Stroop effect decreased with increasing RTs post-incongruent trials. Meanwhile, the slope of the delta plot for cI vs. cC conditions (i.e., ΔCIvCC) was positive (mean= 11.16ms), indicating that the Stroop effect following post-congruent trials increased with increasing RTs. In order to assess the presence of an inflection point in the delta plots, we performed a post-hoc analysis to compare the slope of the final delta plot segment (bins 4-5) with the slope of the early delta plot segments (bins 1-4) within each delta plot using paired t-tests (based on Figure 3). In the ΔIIvIC condition, there was no significant slope difference between early and late RT bins, t(31)=-0.72, p= 0.479, p_adj_= 0.957, suggesting no evidence of an inflection in this condition. In contrast, a significant slope difference between early and late RT bins was observed in the ΔCIvCC condition, t(31)= 2.54, p= 0.016, p_adj_= 0.033).

**Figure 3.**
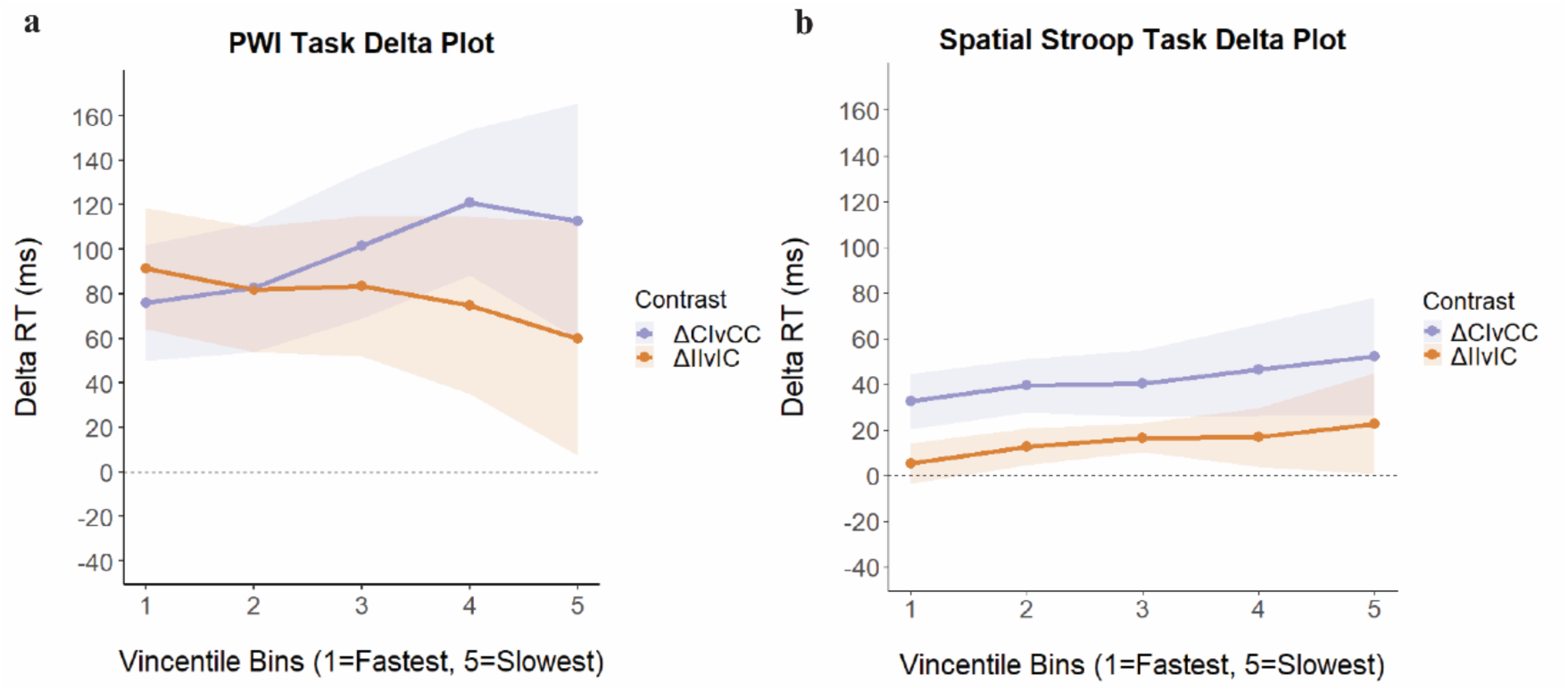
Delta plots between conditions of interest measuring the Stroop Effect in the PWI (a) and spatial Stroop task (b). The x-axis reflects binned reaction times ranging into quintiles from 1-fastest to 5-slowest divided. The y-axis corresponds to the difference in binned response time between contrasted conditions. Shaded regions correspond to 95% confidence intervals. Statistical analysis was performed on subject-level slopes.

**Figure 4.**
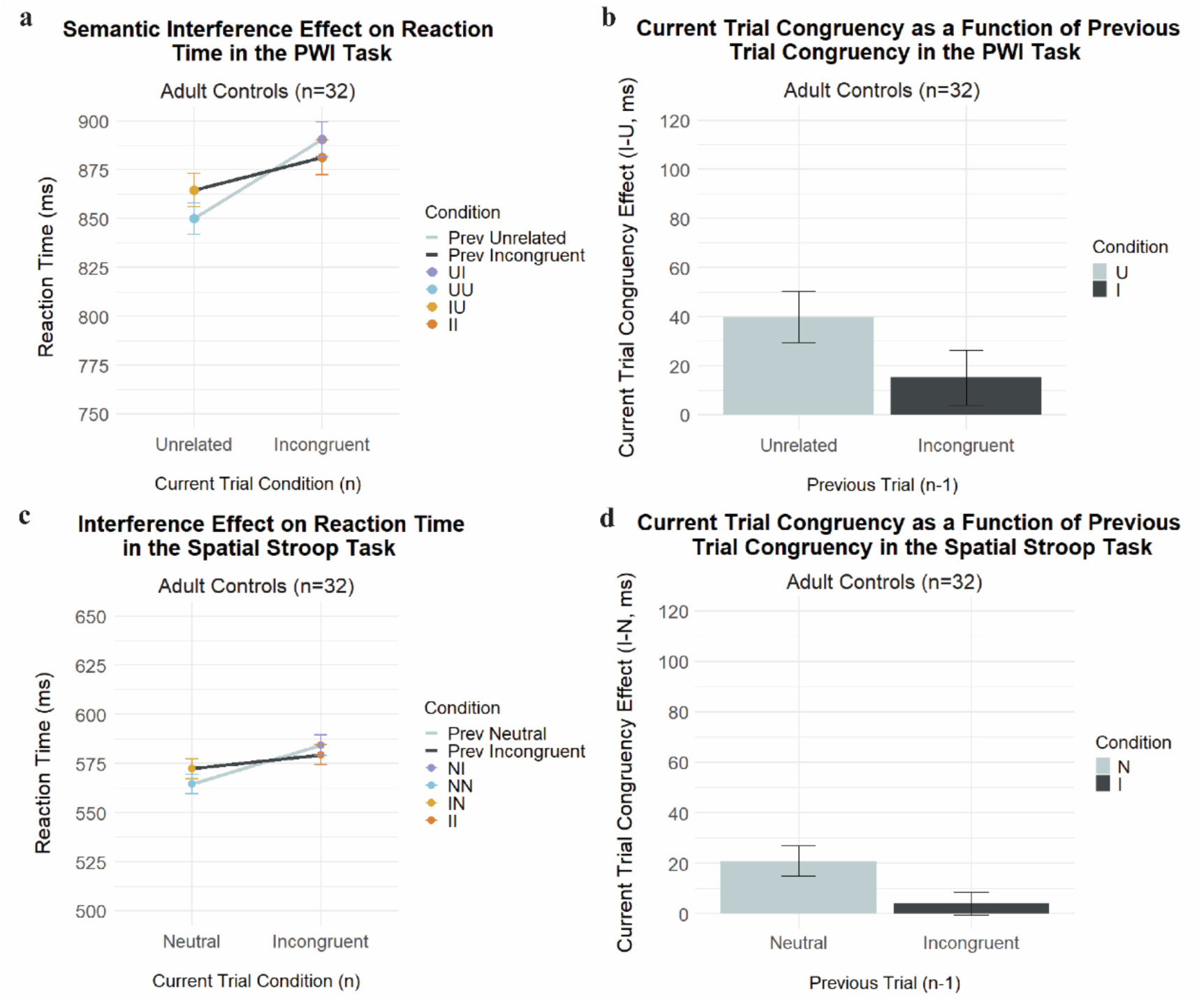
*Left-* CSE effect on reaction times in adults controls in the PWI (a) and Spatial Stroop (c) task in four conditions (i.e., uU-blue, uI-purple, iU-yellow, iI-orange). Previous unrelated trials are the cooler colors (blue and purple) and are connected by a light grey solid line. Previous incongruent trials are the warmer colors (yellow and orange) and are connected by a dark grey solid line. *Right-* Difference between mean RTs in incongruent versus unrelated/neutral trials as a function of previous trial condition (neutral/unrelated, incongruent), illustrating a reduced interference effect following incongruent trials relative to unrelated/neutral trials) in the PWI (b) and spatial Stroop (d) task.

#### 3.1.6. Spatial Stroop Task-Delta Slope Analysis

A repeated-measures ANOVA revealed no significant main effect of condition on delta slopes*, F*(1, 31)= 0.037, *p*= 0.849, *η*p²=.001. The slopes in both conditions were positive (ΔIIvIC: mean = 3.89; ΔCIvCC mean = 4.62), indicating that the Stroop effect increased along the RT distribution when trial n was preceded by a congruent or an incongruent trial. A post-hoc analysis using paired t-tests indicated no significant difference between delta slopes for RT bins 1-4 vs. 4-5 in the ΔIIvIC, t(31)= 1.19, p= 0.243, p_adj_= 0.486, nor in the ΔCIvCC conditions, t(31)=-0.43, p= 0.668, p_adj_= 1.0, indicating that the inflection point found in the PWI task in the ΔCIvCC was not present in the spatial Stroop task.

### 3.2. Part 2: Interference Effect

#### 3.2.1. Interference Effect on Accuracy in PWI and Spatial Stroop

There was a significant main effect of task on Accuracy, χ²(1)= 3.97, p= 0.047, indicating that participants were more accurate in the spatial Stroop task compared to the PWI task (β_raw_=-0.56, SE= 0.28, t=-1.99, p= 0.046). There was no significant main effect of current trial interference (i.e., no difference between incongruent and neutral or unrelated trials, χ²(1)= 3.26, p= 0.071, nor significant main effect of previous trial congruency, χ²(1)= 2.11, p= 0.147, but there was a significant interaction between previous trial interference and current trial interference, χ²(1) = 7.97, p= 0.005, indicating that interference (i.e. difference between incongruent ‘related’ trials and neutral trials) was reduced following incongruent trials. Additionally, there was no significant interaction between previous trial congruency and task, χ²(1)= 1.18, p= 0.278, current trial congruency and task, χ²(1)= 1.31, p= 0.252, nor a three-way interaction between previous trial congruency, current trial congruency and task, χ²(1)= 2.80, p= 0.094, on Accuracy.

#### 3.2.2. Interference Effect on Reaction Times in PWI and Spatial Stroop

There was a significant main effect of Task, χ²(1)= 113.35, p< 0.001. Participants were significantly faster in the spatial Stroop task compared to the PWI task (β_raw_= 2.12×10^-1^, SE=1.99×10^-2^, t= 10.65, p< 0.001). There was no significant main effect of previous trial interference, χ²(1)= 3.45, p= 0.063, nor significant main effect of current trial interference, χ²(1)= 0.47, p= 0.494. There was a significant interaction between previous trial interference and current trial interference, χ²(1)= 11.84, p< 0.001, indicating that the current trial interference effect was reduced following incongruent trials (β_raw_=-9.94×10^-3^, SE= 2.88×10^-1^, t=-3.44, p= 0.009). There was a significant interaction between current trial interference and task, χ²(1)= 7.06, p= 0.008, and the previous trial interference and task, χ²(1)= 4.92, p= 0.026, indicating that the interference effect of the current trial and the previous trial were larger in the PWI than in the spatial Stroop tasks (β_raw_= 9.11×10^-3^, SE= 3.43×10^-3^, t= 2.66, p= 0.008; β_raw_=-6.29×10^-3^, SE= 2.84×10^-3^, t=-2.19, p= 0.027). There was no significant three-way interaction between previous trial interference, current trial interference, and task, χ²(1)= 0.42, p= 0.516, indicating that the modulation of the interference effect by previous trial interference did not significantly differ between the PWI and the spatial Stroop task.

#### 3.2.3. PWI Task-Vincentile Analysis

A repeated-measures ANOVA revealed a significant main effect of current trial congruency, *F*(1, 31)= 8.31, *p*= 0.007, *η*p²= 0.21. As expected, there was a significant main effect of bin, *F*(1, 31)= 501.2, *p*< 0.001, *η*p² =.82. Additionally, there was a significant interaction between current trial congruency and previous trial congruency. F(1,31)= 6.13, *p*= 0.019, *η*p²= 0.17, and a significant three-way interaction between previous trial congruency, current trial congruency, and bin, *F*(1, 31)= 5.42, *p*= 0.027, *η*p²= 0.01, indicating that the magnitude of the sequential semantic interference effect increased as a function of response time. (Figure 5). There was no significant main effect of previous trial congruency, *F*(1, 31) = 0.04, *p*= 0.853, *η*p² =.001, and no significant interaction between current trial congruency and bin, *F*(1, 31)= 0.01, *p*= 0.908, *η*p²<.001, or previous trial congruency and bin, *F*(1, 31)= 0.06, *p*= 0.817, *η*p²<.001.

**Figure 5.**
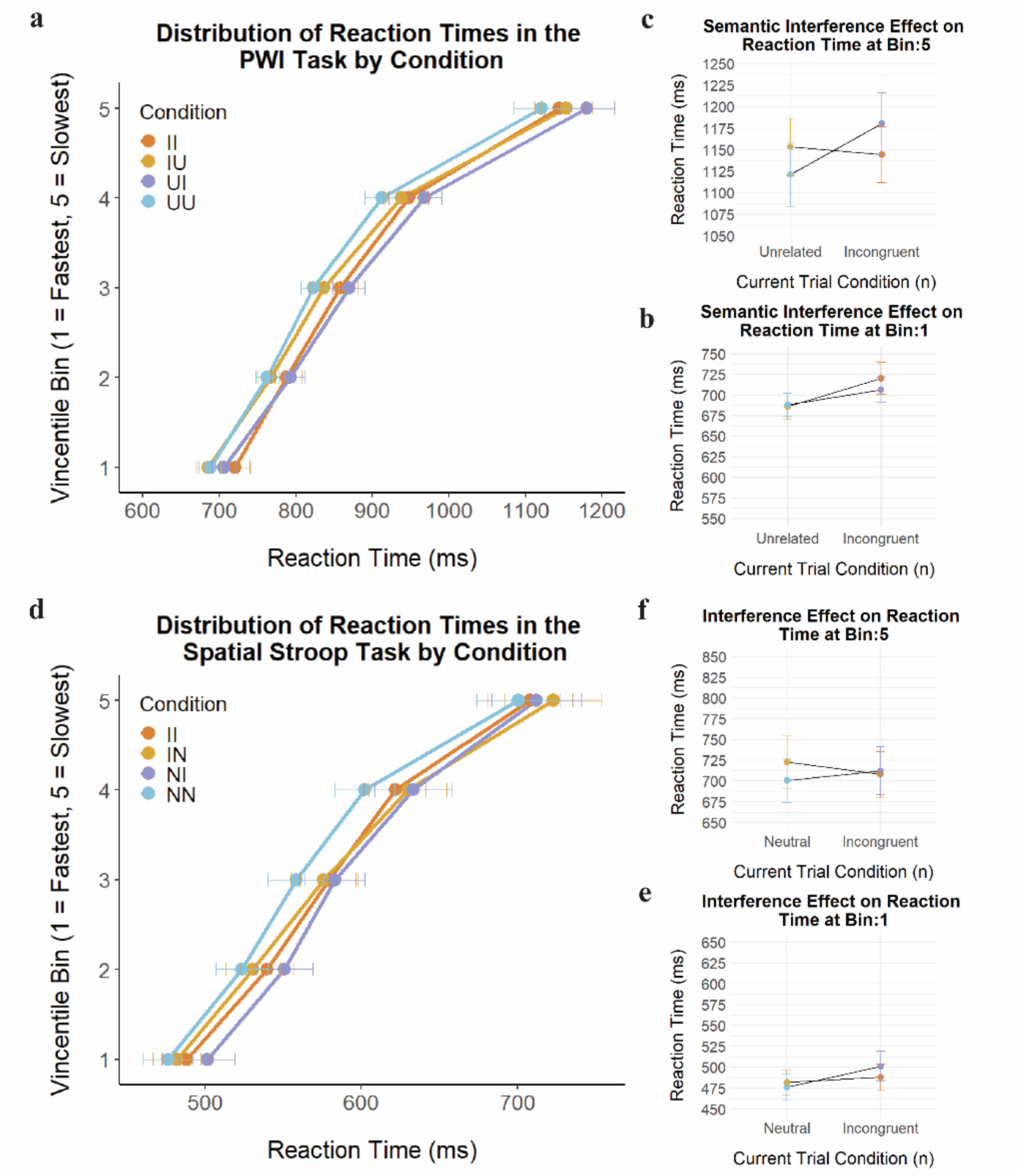
*Left-* Vincentile plots demonstrating the distribution of reaction times across quintile (1-fastest, 5-slowest) by condition (i.e., uU-blue, uI-purple, iU-yellow, iI-orange) in the PWI task (a) and spatial Stroop task (d). Error bars denote the standard error around the mean by condition and bin. *Right-*SI on reaction time at bin 1 and bin 5 in the PWI (top) and spatial Stroop task (bottom). Plots highlight the evolution of the interference effect across the RT distribution from bin 1 (b, e) to 5 (e, f).

**Figure 6.**
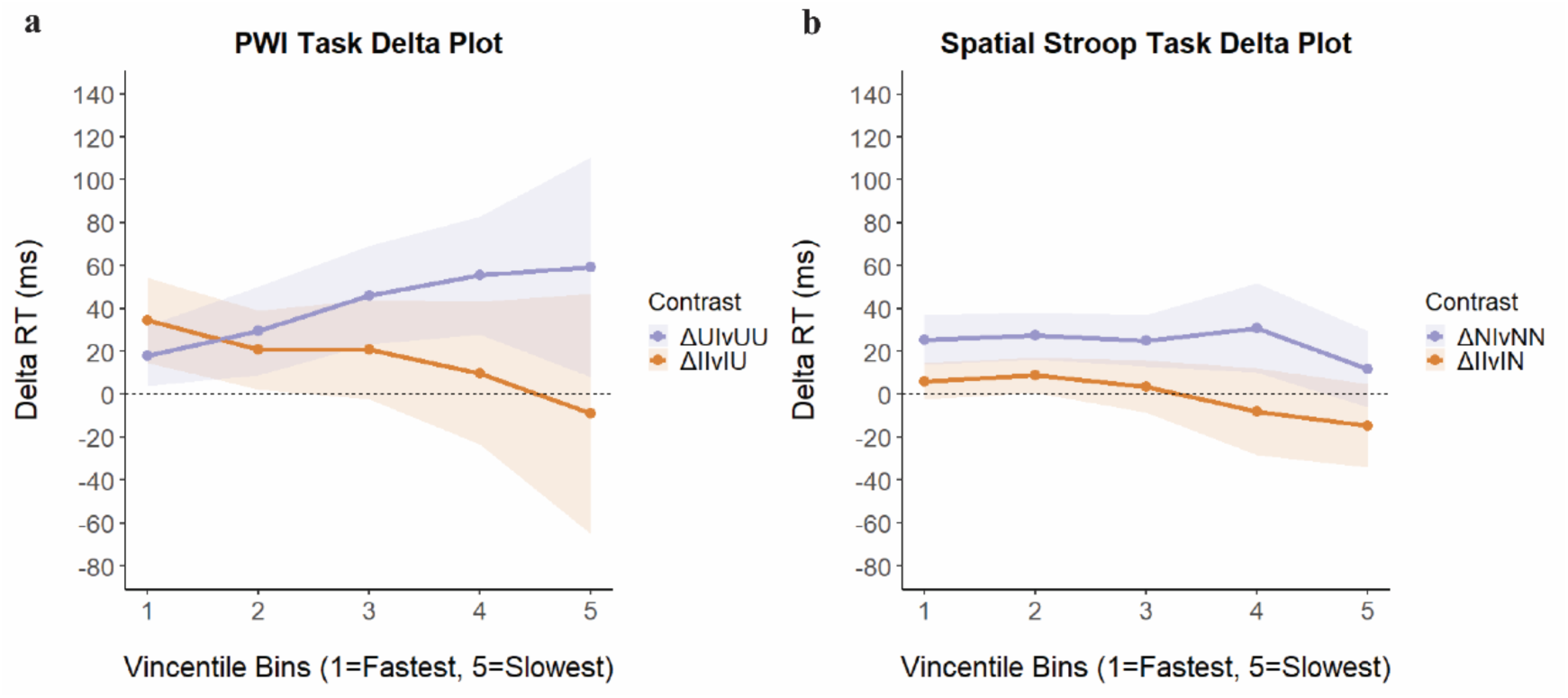
Delta plots of conditions of interest measuring the Interference Effect in the PWI (a) and spatial Stroop task (b). The x-axis reflects binned reaction times ranging from 1-fastest to 5-slowest divided into quintiles. The y-axis corresponds to the difference in binned response time between contrasted conditions. Shaded regions correspond to 95% confidence intervals. Statistical analysis was performed on subject-level slopes.

**Figure 7.**
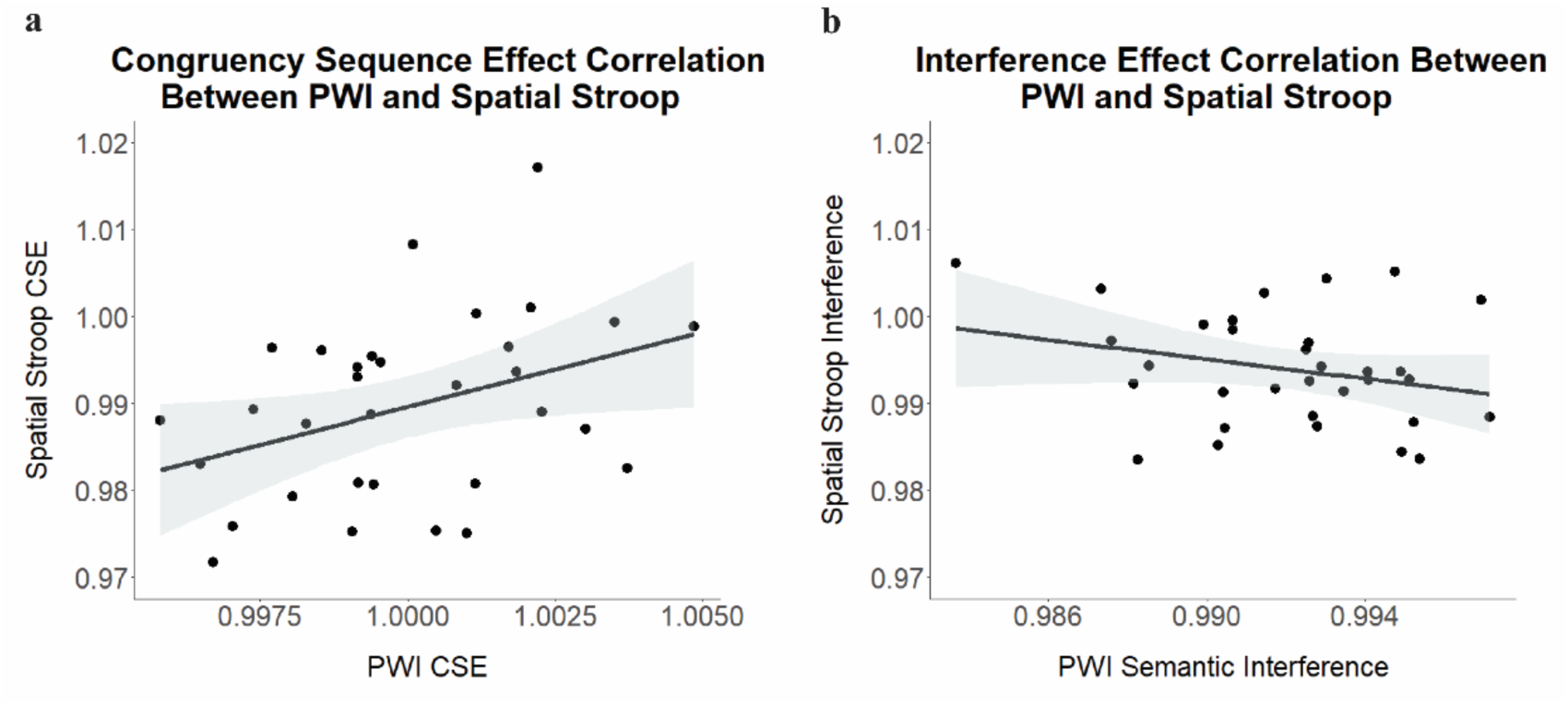
Correlations between participant-specific (a) congruency sequence effects and (b) interference effects exponentiated coefficients between the PWI task and spatial Stroop task.

#### 3.2.4. Spatial Stroop Task-Vincentile Analysis

A repeated-measures ANOVA revealed a significant main effect of current trial congruency (*F*(1, 31) = 11.97, *p* =.002, *η*p² =.28). As expected, there was a significant main effect of quintile (*F*(1, 508) = 2004.56, *p* <.001, *η*p² =.80). There was no significant main effect of previous trial (*F*(1, 31) = 1.06, *p*=.311, *η*p² =.033) and no significant interaction between current trial congruency and quintile (F(1,31)=2.99, *p*=.085, *η*p² =.006), previous trial congruency and quintile (*F*(1, 508) =2.21, *p*=.138, *η*p²=.004), and no significant three-way interaction between previous trial, current trial, and quintile (*F*(1, 508) =.53, *p* =.468, *η*p² =.001).

#### 3.2.5. PWI Task-Slope Analysis

We calculated the difference (delta, Δ) between conditions within each quintile (i.e. Interference effect: ΔIIvIU: iI – iU, ΔUIvUU uI-uU). A repeated-measures ANOVA revealed a significant main effect of condition on slope*, F*(1, 31) = 5.42, *p*= 0.027, *η*p²= 0.15. The slope of the difference between iI and iU conditions (i.e., ΔIIvIU) was negative (mean=-9.78 ms), indicating that the semantic interference effect post-incongruent trials decreased along the RT distribution. Meanwhile, the slope of the difference between uI and uU conditions (i.e. ΔUIvUU) was positive (mean=10.87 ms), indicating that the difference increased post-unrelated trials. A post-hoc analysis using paired t-tests indicated no significant difference in slopes in the ΔIIvIU condition t(31)=-66, p= 0.514, nor in the ΔUIvUU condition, t(31)= 1.14, p= 0.262, between the final delta segment and earlier segments.

#### 3.2.6. Spatial Stroop Task-Slope Analysis

A repeated-measures ANOVA demonstrated no significant main effect of condition on slope*, F*(1, 31)= 0.886, *p*= 0.354, *η*p²= 0.03. The slopes of the difference between iI and iN conditions (i.e., ΔIIvIN) and the difference between nI and nN conditions (i.e., ΔNIvNN) were both slightly negative (mean =-5.86 ms; mean =-2.38 ms). A post-hoc analysis using paired t-tests indicated no significant difference between the slopes of early and late segments in the ΔIIvIU, t(31)= 0.49, p= 0.630, p_adj_= 1.0, nor in the ΔUIvUU, t(31)= 1.67, p= 0.106, p_adj_= 0.216.

### 3.3. Between-Task Correlations

We examined the correlations between sequential congruency and interference effects in the PWI task and the spatial Stroop task. By-participant congruency sequence effects in the linguistic and minimally-linguistic tasks were positively correlated and indicated a small correlation between tasks, *r*(30)= 0.38, t= 2.22, p= 0.034, p_adj_= 0.034). This indicated that participants who showed larger congruency sequence effects in the spatial Stroop task also tended to show larger congruency sequence effects in the PWI task. In contrast, interference effects (i.e., unrelated ‘neutral’ conditions compared to incongruent conditions) were not significantly correlated between tasks, *r*(30)=-0.26, t=-1.49, p= 0.146, p_adj_= 0.146).

## 4. DISCUSSION

Effective communication requires cognitive control. The present study adopted the Dual Mechanisms of Cognitive Control framework (Braver, 2012) to examine the contribution of different types of inhibitory control processes in the resolution of semantic interference in word production. We used a linguistic naming task and a minimally-linguistic inhibitory control task, previously used in bilingualism studies comparing linguistic to non-linguistic conflict resolution (Andrade et al., 2025, Gálvez-mcDonough et al., 2025, Blumenfeld & Marian, 2014, Freeman et al., 2017, Giezen et al., 2015). We analyzed Stroop effects (i.e., incongruent/‘related’ vs congruent), interference effects (i.e., incongruent/‘related’ vs unrelated/neutral), trial-by-trial sequence effects on reaction times and accuracy rates, and measured how these effects varied across the RT distributions. Our findings revealed reliable congruency sequence effects in both the PWI and spatial Stroop tasks. Distributional analyses of reaction times further revealed a dissociation in the way conflict is resolved following a high conflict trial in the PWI compared to the spatial Stroop task. In particular, following semantically-related trials, the Stroop effect decreased with increasing RTs, likely indicating the recruitment of proactive control. Following congruent or neutral trials, the Stroop effect increased with increasing RTs, likely indicating the recruitment of reactive control.

Replicating previous studies (Freund et al., 2016, Shitova et al., 2017), the congruency sequence effect was observed in the PWI task on reaction times. We did not find a Stroop effect on accuracy as seen in Shitova et al., 2017, which may have been due to differences in study design. Specifically, the Shitova et al., 2017 study contained more trials than the present. Additionally, the interference sequence effect on RTs was replicated (as seen in Van Maanen and Van Rijn, 2010, Korko et al., 2021). In the PWI task, this effect manifested as a reduced semantic interference effect after a previous semantically-related trial on reaction times and accuracy rates. Similarly in the spatial Stroop task, there was a reduced incongruent vs. neutral trial difference following incongruent trials. The congruency sequence effect has received much interest in the study of cognitive control processes (Egner, 2007). Importantly, congruency sequence effects have been proposed to stem from different possible sources including conflict-driven adjustments in cognitive control, but also memory effects of stimulus-response associations, as well as repetition expectancy (Egner, 2007). As described in Braem et al. (2019), it can be difficult to dissociate between these possible interpretations in a given paradigm, especially when paradigms include lower level confounds.

Reaction time distributional analyses were used here to attempt to dissociate between the nature of cognitive control processes engaged dynamically in each task. These distributional analyses examine how cognitive inhibitory control mechanisms unfold over time, revealing *when* conflict is preferentially resolved. Previous research using distributional analysis to investigate the semantic interference effect in the PWI task have shown the effect to be *increasing* with inceasing RTs; with some studies showing this effect throughout the RT distribution (Roelofs & Piai, 2017; Fuhrmeister & Bueurki, 2021), and others focusing only on longer RTs (Scaltritti et al., 2015). Increasing effects with increasing RTs are also what we found when only focused on within-trial congruency (part 1) and interference (part 2) effects. In fact, in many tasks eliciting conflict, the Stroop effect has been shown to increase with increasing reaction times (Tang et al., 2022). This has been interpreted as being caused by the prepotent task-irrelevant attribute becoming more active with time within a trial, increasing the time needed for the decision process to happen (Miller and Schwarz, 2021). By teasing apart the trials that were preceded by a high versus low conflict trial, we were able to observe that the pattern of increasing congruency and interference effects with increasing RTs is, in fact, not ubiquitous. Indeed, when the previous trial was semantically-related in the PWI task, interference was largest early in the reaction time distribution and was progressively reduced for longer RTs. Previous studies have associated these variations in slope to differences in activation and suppression mechanisms (Ridderinkhof et al., 2002), with positive-delta slopes indicative of inhibitory processes requiring time to build up to resolve conflict, and negative-delta slopes reflecting active suppression triggered previously to the start of the current trial. Specifically, a postive delta slope following congruent trials is consistent with relatively low state of engagement of adaptive control, resulting in the Stroop effect increasing over time across slower reaction times, until inhibitory control has had sufficient time to accumulate and reduce interference in the slower responses (Ridderinkof et al., 2002). In contrast, a more regulatory, proactive mode of control is observed following an incongruent trial. In this case, control is upregulated, leading to improved performance on subsequent conflict trials. In effect, this upregulation of adaptive control leads to interference effects decreasing over the reaction time distribution, resulting in negative-going delta slopes. In other words, negative-going delta plots can be interpreted as reflecting proactive control triggered by previous high-conflict situations, whereas positive-going delta plots are better aligned with reactive control following a low-conflict situation.

In addition, we observed an inflection point in the delta plots corresponding to the trials following low conflict trials. This inflection point has received interest in prior studies as it has been interpreted as representing the point where inhibitory control has had enough time to accumulate following the build-up of conflict in the current trial. Observing the delta plot dissociation and inflection point we observe in the PWI task indicates that reactive and proactive control are dynamically engaged to resolve conflict in word production. This dissociation was not found in the spatial Stroop task eventhough there were minimal differences between tasks in the CSE and the size of the mean congruency effects were found to correlate across participants.

Observing the delta plot dissociation in the PWI task indicates that reactive and proactive control are dynamically engaged to resolve conflict in word production. This dissociation was not found in the spatial Stroop task eventhough there were minimal differences between tasks in the CSE and the size of the mean Stroop effects were found to correlate across participants. One way to situate this difference in delta plot patterns between the PWI and spatial Stroop paradigms within current theoretical frameworks is by understanding the nature of the conflict elicited in each task. As noted in the introduction, both paradigms are proposed to induce prepotent conflict, where the dominant but task-irrelvant features (i.e., the distractor word or arrow location) must be overriden, in favor of a goal-directed response (i.e., naming the picture or the direction the arrow is pointing towards). However, there are important differences between the two paradigms. In the spatial Stroop task, the stimulus set is much smaller than in the PWI task as there is only a limited number of directions (i.e., left and right) and locations the arrow can be presented in. The reduced stimulus options available can therefore lead to memory effects compared to studies with greater stimulus variation (Braem et al., 2019, Mayr et al., 2003, Hommel Hommel et al., 2004). Because of the small stimulus set, the previous incongruent trial can belong to only one of two alternatives, resulting in the consecutive presentation of identical stimuli in the spatial Stroop task. This leads to a feature confound as the trials that induce subsequent control adjustments (i.e., inducer trials) are identical to the trials where these control adjustments are measured (i.e., diagnostic trials according to the Braem et al., 2019 framework). This is not the case in the PWI task given the much larger stimulus set and the fact that the same picture or distractor words were never displayed twice in a row. According to Braem et al., 2019, this dissociation between inducer and diagnostic trials is essential to capturing true adaptive control processes that cannot be as easily confounded by low level feature expectations. From this perspective, the PWI task is much better suited to capture conflict adaptation from conflict-based learning (Braem, 2019, Duthroo et al., 2014) than the spatial Stroop task, even though the spatial Stroop task has been used extensively to compare linguistic to non-linguistic processing (Andrade et al., 2025, Gálvez-mcDonough et al., 2025, Freeman et al., 2017, Giezen et al., 2015). Importantly, the same argument can be made for other studies comparing linguistic tasks with rich stimulus sets to non-linguistic cognitive control paradigms involving restricted stimulus sets, such as the Flanker task (as in e.g. Mendoza et al., 2021), the Simon task (e.g., Riès et al., 2013), etc. The limited stimulus set could explain why no dissociation or inflection point were observed in the spatial Stroop paradigm. Instead, small within and between trial congruency effects were observed in this task along with a weak positive build-up of conflict across the RT distribution, consistent with a relatively low state engagement of inhibitory control. Importantly, this difference in the engagement of pro-versus reactive control mechanisms between paradigms would not have been visible without the fine-grained distributional analyses performed and indicate that comparing control processes involved in and outside language requires more thoughtful considerations regarding experimental design.

## 5. CONCLUSION

Language production involves cognitive control. Using the dual mechanisms of cognitive control framework (Braver, 2012) and distributional reaction time analyses (Luce, 1991, Vincent, 1912, De Jong et al., 1994), we found that proactive and reactive control are dynamically engaged to support language production. In particular, following high conflict situations, proactive control acts as a top-down anticipatory mechanism to resolve within-language conflict before a word is even spoken, minimizing interruptions when conversing while reactive control is still recruited in an adaptive way to resolve conflict following low conflict situations. Importantly, comparing linguistic conflict resolution to non or minimally-linguistic conflict resolution revealed that the paradigms used must be carefully matched beyond simply involving the need to overcome a prepotent response. This finding has important implications for studies investigating parallels between language and other domains given the inherent vast repertoire of representations present in language. While exact matching of stimuli sets may be difficult to accomplish, the principles developed in the broad field of cognitive control may still be instrumental in the understanding of the underlying processes of language production. In addition, while proactive and reactive control in the linguistic domain have been largly explored through the lens of bilingualism to resolve interference between languages, our study illustrates that proactive and reactive control mechanisms are also present when resolving interference within a language. In addition, while deficits in proactive control have been observed in populations with neuropsychological disorders (Wylie et al., 2009, Clawson et al., 2013, Abrahamse, 2016, Kricheldorff et al., 2023) and aging (Ileri-Taylor et al., 2025), relatively little is known about the roles of the dual modes of cognitive control in neurotypical language production. Understanding how these control mechanisms contribute to language production is essential to clarify how speakers plan, monitor, and adapt their utterances in real time.

## SUPPLEMENTARY MATERIALS

### S1: Neuropsychological Assessments

Participants were administered the Montreal Cognitive Assessment (MOCA; Nasreddine et al., 2005) to assess cognitive function, the Boston Naming Task (BNT; Kaplan et al., 1983) to assess naming ability, and the Test of Premorbid Functioning (TOPF; Pearson Clinical, 2017) to assess single word reading abilities. On average, cognitive scores fell within the normal range based on published norms for the MOCA and TOPF. On the BNT, 71.87% of participant’s scores fell within the borderline to low average range, whereas 28.13% of scores fell within the average to superior range. Prior research has demonstrated group differences in BNT performance, with African Americans, Asians, and Hispanics individuals-who comprise 56% of the present sample size-showing lower performance compared to Caucasian Americans (Boone et al., 2007,Pedraza et al., 2009, Strauss et al., 2006, Whitfield et al., 2000). Additionally, other contextual factors such as geographic location and socio-economic status can influence the frequency with which objects assessed by the BNT are encountered in daily life. Limited exposure to certain test items may therefore contribute to the naming errors that are independent from underlying neurological pathology (Barker-Collo, 2001, Beattey et al., 2017, Hamberger & Seidel, 2003). Collectively, these factors may account for the relatively lower BNT performance in this sample, despite overall high average naming performance across participants (see section 3).

### S2: Follow up Delta plot analysis on younger adults

Data from older participants were removed to restrict the sample to young adults, given established age-related differences in cognitive control (Weissberger et al., 2012, de Bruin et al., 2020, de Bruin et al., 2023). Therefore, analysis was performed on the remaining 27 participants (20 females, mean age: 26.33 years, SD: 4.93, mean education: 16.7 years, SD: 1.7).

*Stroop Effect*: For each participant, we calculated the difference (delta Δ) between conditions within each bin (i.e. ΔIIvIC: iI – iC, ΔCIvCC: cI – cC). In the PWI task, a repeated-measures ANOVA revealed a significant main effect of Condition on slope*, F*(1, 26)= 5.77, *p*= 0.024, *η*p²= 0.18. The slope of the delta plot for iI vs. iC conditions (i.e., ΔIIvIC) was negative (mean=-11.58 ms), indicating that the congruency effect decreased with increasing RTs post-incongruent trials. Meanwhile, the slope of the difference between cI and cC conditions (i.e., ΔCIvCC) was positive (mean= 9.38), indicating that the congruency effect following post-congruent trials increased with increasing RTs. In the spatial Stroop task, no significant main effect of Condition on slope was present*, F*(1,26) = 0.041, *p*= 0.842, *η*p²= 0.002. The slopes of the difference between iI and iC conditions (i.e., ΔIIvIC) and the difference between cI and cC conditions (i.e., ΔCIvCC) were both positive (mean = 2.61 ms; mean = 3.49 ms, indicating that the congruency effect increased along the RT distribution when trial n was preceded by a congruent or an incongruent trial.

Interference Effect: We calculated the difference (delta Δ) between conditions within each quintile (ΔIIvIU: iI – iU, ΔUIvUU uI-uU). In the PWI task, a repeated-measures ANOVA revealed no significant main effect of Condition on slope*, F*(1,26)= 2.98, *p*=.096, *η*p²= 0.10. The slope of the difference between iI and iU conditions (i.e., ΔIIvIU) was more negative (mean=-8.02 ms) relative to the slope of the difference between uI and uU conditions (i.e., ΔUIvUU; mean= 9.76 ms) condition. In the spatial Stroop task, no significant main effect of Condition on slope was present*, F*(1,26) = 0.04, *p*= 0.843, *η*p²= 0.002. The slope of the difference between iI and iN conditions (i.e., ΔIIvIU) and the difference between nI and nN conditions (i.e., ΔNIvNN) were both negative (mean=-4.63 ms; mean= - 3.88 ms).

^i^ A limitation of this study was the broad age range of participants, recruited in part to align with a co-occurring study. Prior bilingual language-control research suggests that proactive and reactive control processes may vary with age, particularly in older adults (>60 years) (e.g., Weissberger et al., 2012; de Bruin et al., 2020, 2023). However, our sample consisted primarily of young and middle-aged adults (18–59 years), among whom proactive control remains detectable, though diminished (Ileri-Tayar et al., 2025). Follow-up analyses excluding participants over 40 years (see Supplementary Materials) yielded attenuated effects, likely reflecting reduced statistical power. Future work should more directly examine the role of age and language experience in proactive and reactive control mechanisms.

---

**Figure S2.**
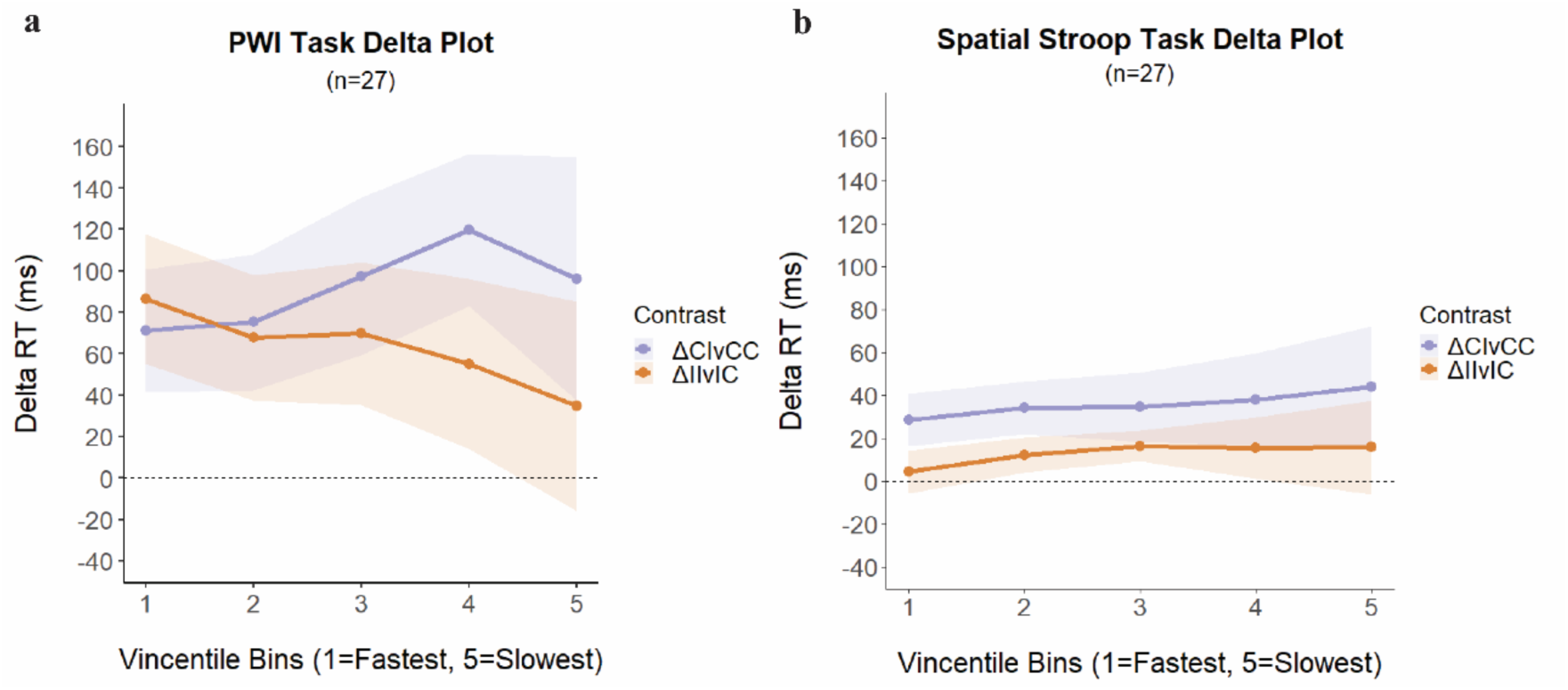
Delta plots between conditions of interest measuring the Stroop Effect in the PWI (a) and spatial Stroop task (b) in young adults. The x-axis reflects binned reaction times ranging from 1-fastest to 5-slowest divided into quintiles. The y-axis corresponds to the difference in binned response time between contrasted conditions. Shaded regions correspond to 95% confidence intervals.

**Figure S3.**
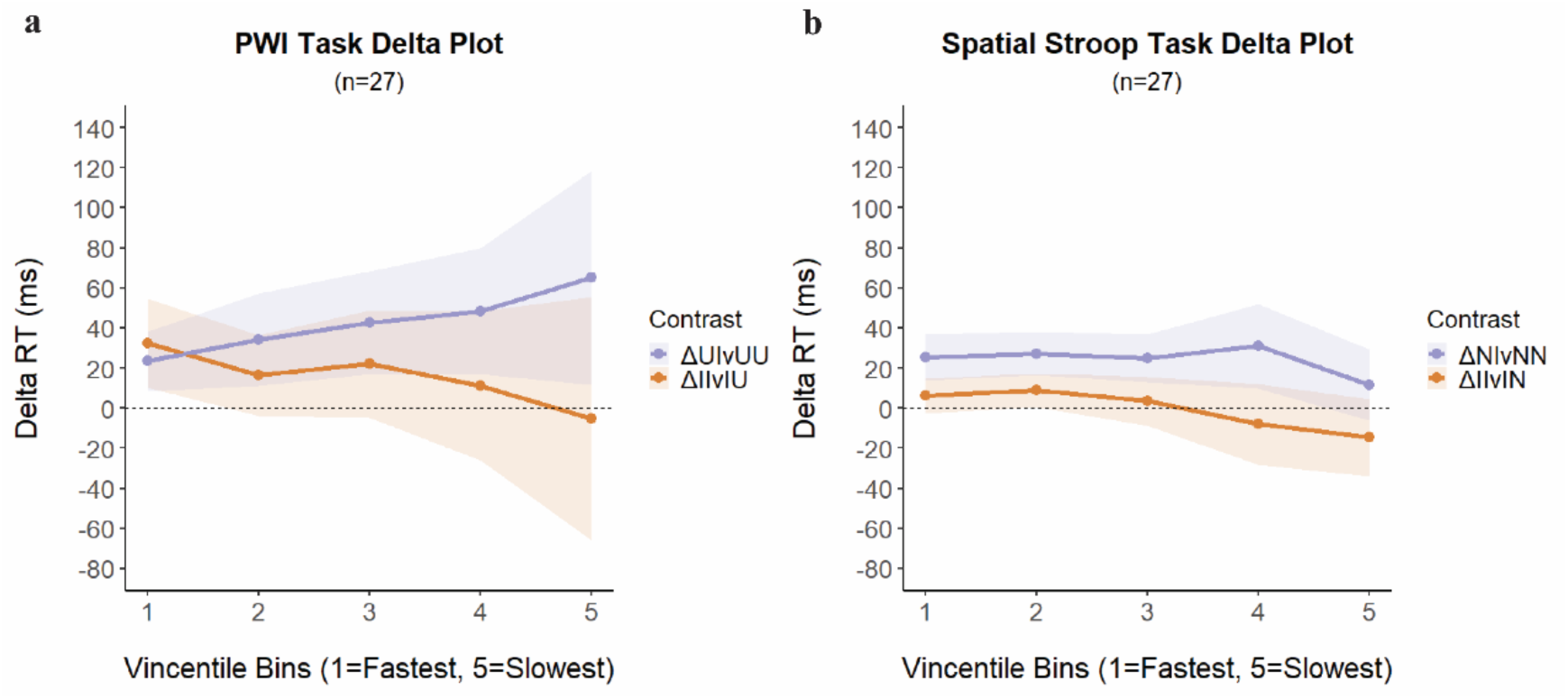
Delta plots of conditions of interest measuring the Interference Effect in the PWI (a) and spatial Stroop task (b) in young adults. The x-axis reflects binned reaction times ranging from 1-fastest to 5-slowest divided into quintiles. The y-axis corresponds to the difference in binned response time between contrasted conditions. Shaded regions correspond to 95% confidence intervals.

**Table S1:** Neuropsychological assessment: Montreal Cognitive Assessment (MOCA); Boston Naming Task (BNT), and the Test of Premorbid Functioning (TOPF). Note: BNT raw scores are from spontaneous naming. Additionally, MOCA ‘percentile rank’ scores denote percentage correct as opposed to traditionally percentile rank that is compared to a normative group.

| *N*=32 | Raw Score: | Percentile Rank: | Scaled Score: |
| --- | --- | --- | --- |
|  | Mean (SD; range) | Mean (SD; range) | Mean (SD; range) |
| MOCA | 28.03 (1.6; 26-30) | 93.43 (5.2; 86-100) | - |
| TOPF | 51 (10.2; 33-66) | 72.06 (19.6; 34-97) | 110.47 (9.6; 94-128) |
| BNT | 53.47 (3.67; 47-60) | 27.56 (32.65; 3-92) | 9.41 (2.8; 7-16) |

## REFERENCES

Acheson, D. J., & MacDonald, M. C. (2009). Verbal working memory and language production: Common approaches to the serial ordering of verbal information. Psychological Bulletin, 135(1), 50–68. 10.1037/a0014411

Andrade, K. D., Blumenfeld, H. K., & Riès, S. K. (2025). Brain dynamics of crosslinguistic interference resolution in Spanish–English bilinguals with and without aphasia. Bilingualism: Language and Cognition, 1–19. 10.1017/S1366728925100461

Baayen, R. H., Davidson, D. J., & Bates, D. M. (2008). Mixed-effects modeling with crossed random effects for subjects and items. *Journal of Memory and Language*, Special Issue: Emerging Data Analysis, 59(4), 390–412. 10.1016/j.jml.2007.12.005

Balota, D., & Yap, M. (2011). Moving Beyond the Mean in Studies of Mental Chronometry: The Power of Response Time Distributional Analyses. Current Directions in Psychological Science - CURR DIRECTIONS PSYCHOL SCI, 20, 160–166. 10.1177/0963721411408885

Barker-Collo, S. L. (2001). The 60-Item Boston Naming Test: Cultural bias and possible adaptations for New Zealand. Aphasiology, 15(1), 85–92. 10.1080/02687040042000124

Beattey, R. A., Murphy, H., Cornwell, M., Braun, T., Stein, V., Goldstein, M., & Bender, H. A. (2017). Caution warranted in extrapolating from Boston Naming Test item gradation construct. Applied Neuropsychology. Adult, 24(1), 65–72. 10.1080/23279095.2015.1089505

Belke, E., Meyer, A. S., & Damian, M. F. (2005). Refractory effects in picture naming as assessed in a semantic blocking paradigm. The Quarterly Journal of Experimental Psychology A: Human Experimental Psychology, *58A*(4), 667–692. 10.1080/02724980443000142

Blumenfeld, H. K., & Marian, V. (2014). Cognitive control in bilinguals: Advantages in Stimulus-Stimulus inhibition. Bilingualism, 17(3), 610–629. 10.1017/S1366728913000564

Boone, K. B., Victor, T. L., Wen, J., Razani, J., & Pontón, M. (2007). The association between neuropsychological scores and ethnicity, language, and acculturation variables in a large patient population. Archives of Clinical Neuropsychology: The Official Journal of the National Academy of Neuropsychologists, 22(3), 355–365. 10.1016/j.acn.2007.01.010

Botvinick, M. M., Braver, T. S., Barch, D. M., Carter, C. S., & Cohen, J. D. (2001). Conflict monitoring and cognitive control. Psychological Review, 108(3), 624–652. 10.1037/0033-295x.108.3.624

Braem, S., Bugg, J. M., Schmidt, J. R., Crump, M. J. C., Weissman, D. H., Notebaert, W., & Egner, T. (2019). Measuring Adaptive Control in Conflict Tasks. Trends in Cognitive Sciences, 23(9), 769–783. 10.1016/j.tics.2019.07.002

Braver, T. S. (2012). The variable nature of cognitive control: A dual mechanisms framework. Trends in Cognitive Sciences, 16(2), 106–113. 10.1016/j.tics.2011.12.010

Braver, T. S., Barch, D. M., Keys, B. A., Carter, C. S., Cohen, J. D., Kaye, J. A., Janowsky, J. S., Taylor, S. F., Yesavage, J. A., Mumenthaler, M. S., Jagust, W. J., & Reed, B. R. (2001). Context processing in older adults: Evidence for a theory relating cognitive control to neurobiology in healthy aging. Journal of Experimental Psychology. General, 130(4), 746–763.

Braver, T. S., Gray, J. R., & Burgess, G. C. (2007). Explaining the many varieties of working memory variation: Dual mechanisms of cognitive control. In Variation in working memory (pp. 76–106). Oxford University Press.

Braver, T. S., Gray, J. R., & Burgess, G. C. (2008). Explaining the Many Varieties of Working Memory Variation: Dual Mechanisms of Cognitive Control. In A. Conway, C. Jarrold, M. Kane, A. Miyake, & J. Towse (Eds.), Variation in Working Memory (p. 0). Oxford University Press. 10.1093/acprof:oso/9780195168648.003.0004

Brodeur, M. B., Guérard, K., & Bouras, M. (2014). Bank of Standardized Stimuli (BOSS) phase II: 930 new normative photos. PloS One, 9(9), e106953. 10.1371/journal.pone.0106953

Clawson, A., Clayson, P. E., & Larson, M. J. (2013). Cognitive control adjustments and conflict adaptation in major depressive disorder. Psychophysiology, 50(8), 711–721. 10.1111/psyp.12066

de Bruin, A., Kressel, H., & Hemmings, D. (2023). A comparison of language control while switching within versus between languages in younger and older adults. Scientific Reports, 13(1), 16740. 10.1038/s41598-023-43886-1

de Bruin, A., Samuel, A. G., & Duñabeitia, J. A. (2020). Examining bilingual language switching across the lifespan in cued and voluntary switching contexts. Journal of Experimental Psychology: Human Perception and Performance, 46(8), 759–788. 10.1037/xhp0000746

De Jong, R., Liang, C. C., & Lauber, E. (1994). Conditional and unconditional automaticity: A dual-process model of effects of spatial stimulus-response correspondence. Journal of Experimental Psychology. Human Perception and Performance, 20(4), 731–750. 10.1037//0096-1523.20.4.731

Dell’Acqua, R., Job, R., Peressotti, F., & Pascali, A. (2007). The picture-word interference effect is not a Stroop effect. Psychonomic Bulletin & Review, 14(4), 717–722. 10.3758/BF03196827

Diamond, A. (2013). Executive functions. Annual Review of Psychology, 64, 135–168. 10.1146/annurev-psych-113011-143750

Douglas Bates, Martin Maechler, Ben Bolker, Steve Walker (2015). Fitting Linear Mixed-Effects Models Using lme4. Journal of Statistical Software, 67(1), 1–48. doi:10.18637/jss.v067.i01.

Egner, T. (2007). Congruency sequence effects and cognitive control. *Cognitive, Affective*, & Behavioral Neuroscience, 7(4), 380–390. 10.3758/CABN.7.4.380

Egner, T. (2014). Creatures of habit (and control): A multi-level learning perspective on the modulation of congruency effects. Frontiers in Psychology, 5. 10.3389/fpsyg.2014.01247

Egner, T., Ely, S., & Grinband, J. (2010). Going, Going, Gone: Characterizing the Time-Course of Congruency Sequence Effects. Frontiers in Psychology, 1. 10.3389/fpsyg.2010.00154

Engle, R. W. (2018). Working Memory and Executive Attention: A Revisit. Perspectives on Psychological Science: A Journal of the Association for Psychological Science, 13(2), 190–193. 10.1177/1745691617720478

Eriksen, B. A., & Eriksen, C. W. (1974). Effects of noise letters upon the identification of a target letter in a nonsearch task. Perception & Psychophysics, 16(1), 143–149. 10.3758/BF03203267

Ferreira, V. S., & Pashler, H. (2002). Central bottleneck influences on the processing stages of word production. Journal of Experimental Psychology. Learning, Memory, and Cognition, 28(6), 1187–1199.

Freeman, M. R., Blumenfeld, H. K., & Marian, V. (2017). Cross-linguistic phonotactic competition and cognitive control in bilinguals. Journal of Cognitive Psychology, 29(7), 783–794. 10.1080/20445911.2017.1321553

Freund, M., Gordon, B., & Nozari, N. (2016). Conflict-based regulation of control in language production. Proceedings of the Annual Meeting of the Cognitive Science Society, 38(0). https://escholarship.org/uc/item/59h5p1m1

Freund, M., & Nozari, N. (2018). Is adaptive control in language production mediated by learning? Cognition, 176, 107–130. 10.1016/j.cognition.2018.03.009

Fox J, Weisberg S (2019). _An R Companion to Applied Regression_, Third edition. Sage, Thousand Oaks CA.

Fuhrmeister, P., & Bueurki, A. (2021). *Distributional properties of semantic interference in picture naming: Bayesian meta-analyses* (arXiv:2011.09316). arXiv. 10.48550/arXiv.2011.09316

Fuhrmeister, P., & Bürki, A. (2022). Distributional properties of semantic interference in picture naming: Bayesian meta-analyses. Psychonomic Bulletin & Review, 29(2), 635–647. 10.3758/s13423-021-02016-6

Georgopoulos, A. P. (2002). Cognitive motor control: Spatial and temporal aspects. Current Opinion in Neurobiology, 12(6), 678–683. 10.1016/s0959-4388(02)00382-3

Giezen, M. R., Blumenfeld, H. K., Shook, A., Marian, V., & Emmorey, K. (2015). Parallel language activation and inhibitory control in bimodal bilinguals. Cognition, 141, 9–25. 10.1016/j.cognition.2015.04.009

Gratton, G., Coles, M. G., & Donchin, E. (1992). Optimizing the use of information: Strategic control of activation of responses. Journal of Experimental Psychology. General, 121(4), 480–506. 10.1037//0096-3445.121.4.480

Hamberger, M. J., & Seidel, W. T. (2003). Auditory and visual naming tests: Normative and patient data for accuracy, response time, and tip-of-the-tongue. Journal of the International Neuropsychological Society: JINS, 9(3), 479–489. 10.1017/s135561770393013x

Hommel, B., Proctor, R. W., & Vu, K.-P. L. (2004). A feature-integration account of sequential effects in the Simon task. Psychological Research, 68(1), 1–17. 10.1007/s00426-003-0132-y

Howard, D., Nickels, L., Coltheart, M., & Cole-Virtue, J. (2006). Cumulative semantic inhibition in picture naming: Experimental and computational studies. Cognition, 100(3), 464–482. 10.1016/j.cognition.2005.02.006

Jaeger, T. F. (2008). Categorical data analysis: Away from ANOVAs (transformation or not) and towards logit mixed models. *Journal of Memory and Language*, Special Issue: Emerging Data Analysis, 59(4), 434–446. 10.1016/j.jml.2007.11.007

Jiang, Y., Rouder, J. N., & Speckman, P. L. (2004). A note on the sampling properties of the Vincentizing (quantile averaging) procedure. Journal of Mathematical Psychology, 48(3), 186–195. 10.1016/j.jmp.2004.01.002

Kaplan, E., Goodglass, H., & Weintrab, S. (1983). The Boston naming test. Philadelphia: Lea & Febiger.

Korko, M., Coulson, M., Jones, A., & de Mornay Davies, P. (2021). Types of interference and their resolution in monolingual word production. Acta Psychologica, 214, 103251. 10.1016/j.actpsy.2021.103251

Kornblum, S., Hasbroucq, T., & Osman, A. (1990). Dimensional overlap: Cognitive basis for stimulus-response compatibility--A model and taxonomy. Psychological Review, 97(2), 253–270. 10.1037/0033-295X.97.2.253

Kricheldorff, J., Ficke, J., Debener, S., & Witt, K. (2023). Impaired proactive cognitive control in Parkinson’s disease. Brain Communications, 5(6), fcad327. 10.1093/braincomms/fcad327

Kuznetsova A, Brockhoff PB, Christensen RHB (2017). “lmerTest Package: Tests in Linear Mixed Effects Models.” _Journal of Statistical Software_, *82*(13), 1-26. <10.18637/jss.v082.i13>

Liu, X., Banich, M. T., Jacobson, B. L., & Tanabe, J. L. (2004). Common and distinct neural substrates of attentional control in an integrated Simon and spatial Stroop task as assessed by event-related fMRI. NeuroImage, 22(3), 1097–1106. 10.1016/j.neuroimage.2004.02.033

Luce, R. D. (1991). Response Times: Their Role in Inferring Elementary Mental Organization. Oxford University Press.

Lupker, S. J. (1979). The semantic nature of response competition in the picture-word interference task. Memory & Cognition, 7(6), 485–495. 10.3758/BF03198265

MacLeod, C. M. (1991). Half a century of research on the Stroop effect: An integrative review. Psychological Bulletin, 109(2), 163–203. 10.1037/0033-2909.109.2.163

Matsushima, E. H., & Aznar-Casanova, J. A. (2025). Reconciling the Neurophysiological and Cognitive Theories of Stimulus–Response Spatial Compatibility Effects: A Visual–Motor Dissociation Approach. Vision, 9(2), 34. 10.3390/vision9020034

Mayr, U., Awh, E., & Laurey, P. (2003). Conflict adaptation effects in the absence of executive control. Nature Neuroscience, 6(5), 450–452. 10.1038/nn1051

Meyer, H. C., & Bucci, D. J. (2016). Neural and behavioral mechanisms of proactive and reactive inhibition. Learning & Memory, 23(10), 504–514. 10.1101/lm.040501.115

Miller, E. K., & Cohen, J. D. (2001). An integrative theory of prefrontal cortex function. Annual Review of Neuroscience, 24, 167–202. 10.1146/annurev.neuro.24.1.167

Miller, J., & Schwarz, W. (2021). Delta plots for conflict tasks: An activation-suppression race model. Psychonomic Bulletin & Review, 28(6), 1776–1795. 10.3758/s13423-021-01900-5

Miyake, A., Friedman, N. P., Emerson, M. J., Witzki, A. H., Howerter, A., & Wager, T. D. (2000). The unity and diversity of executive functions and their contributions to complex “Frontal Lobe” tasks: A latent variable analysis. Cognitive Psychology, 41(1), 49–100. 10.1006/cogp.1999.0734

Nasreddine, Z. S., Phillips, N. A., Bédirian, V., Charbonneau, S., Whitehead, V., Collin, I., Cummings, J. L., & Chertkow, H. (2005). The Montreal Cognitive Assessment, MoCA: A brief screening tool for mild cognitive impairment. Journal of the American Geriatrics Society, 53(4), 695–699. 10.1111/j.1532-5415.2005.53221.x

Nozari, N. (2025). The relationship between monitoring, control, conscious awareness and attention in language production. Journal of Neurolinguistics, 74, 101247. 10.1016/j.jneuroling.2025.101247

Nozari, N., & Hepner, C. R. (2019). To select or to wait? The importance of criterion setting in debates of competitive lexical selection. Cognitive Neuropsychology, 36(5–6), 193–207. 10.1080/02643294.2018.1476335

Oppenheim, G. M., Dell, G. S., & Schwartz, M. F. (2010). The dark side of incremental learning: A model of cumulative semantic interference during lexical access in speech production. Cognition, 114(2), 227–252. 10.1016/j.cognition.2009.09.007

Pearson Clinical (2017). TOPF (test of pre-morbid function): Case studies. San Antonio, TX: Pearson Clinical.

Pedraza, O., Graff-Radford, N. R., Smith, G. E., Ivnik, R. J., Willis, F. B., Petersen, R. C., & Lucas, J. A. (2009). Differential Item Functioning of the Boston Naming Test in Cognitively Normal African American and Caucasian Older Adults. Journal of the International Neuropsychological Society: JINS, 15(5), 758–768. 10.1017/S1355617709990361

Piai, V., & Zheng, X. (2019). Speaking waves: Neuronal oscillations in language production. 302. 10.1016/bs.plm.2019.07.002

Protopapas, A. (2007). CheckVocal: A program to facilitate checking the accuracy and response time of vocal responses from DMDX. Behavior Research Methods, 39(4), 859–862. 10.3758/bf03192979

Ratcliff, R. (1979). Group reaction time distributions and an analysis of distribution statistics. Psychological Bulletin, 86(3), 446–461.

Ridderinkhof, K. R. (2002). Micro-and macro-adjustments of task set: Activation and suppression in conflict tasks. Psychological Research, 66(4), 312–323. 10.1007/s00426-002-0104-7

Ridderinkhof, K. R., Ullsperger, M., Crone, E. A., & Nieuwenhuis, S. (2004). The role of the medial frontal cortex in cognitive control. Science, 306(5695), 443–447. 10.1126/science.1100301

Riès, S. K., Xie, K., Haaland, K. Y., Dronkers, N. F., & Knight, R. T. (2013). Role of the lateral prefrontal cortex in speech monitoring. Frontiers in Human Neuroscience, 7, 703. 10.3389/fnhum.2013.00703

Roelofs, A., & Piai, V. (2017). Distributional analysis of semantic interference in picture naming. Quarterly Journal of Experimental Psychology (2006), *70*(4), 782–792. 10.1080/17470218.2016.1165264

Roelofs, A., Piai, V., & Garrido Rodriguez, G. (2011). Attentional Inhibition in Bilingual Naming Performance: Evidence from Delta-Plot Analyses. Frontiers in Psychology, 2. 10.3389/fpsyg.2011.00184

Roquet, A., Poletti, C., & Lemaire, P. (2020). Sequential modulations of executive control processes throughout lifespan in numerosity comparison. Cognitive Development, 54, 100884. 10.1016/j.cogdev.2020.100884

Scaltritti, M., Navarrete, E., & Peressotti, F. (2015). Distributional analyses in the picture–word interference paradigm: Exploring the semantic interference and the distractor frequency effects. Quarterly Journal of Experimental Psychology, 68(7), 1348–1369. 10.1080/17470218.2014.981196

Scherbaum, S., Dshemuchadse, M., Ruge, H., & Goschke, T. (2012). Dynamic goal states: Adjusting cognitive control without conflict monitoring. NeuroImage, 63(1), 126–136. 10.1016/j.neuroimage.2012.06.021

Schriefers, H., Meyer, A. S., & Levelt, W. J. M. (1990). Exploring the time course of lexical access in language production: Picture-word interference studies. Journal of Memory and Language, 29(1), 86–102. 10.1016/0749-596X(90)90011-N

Schwarz, W., & Miller, J. (2012). Response time models of delta plots with negative-going slopes. Psychonomic Bulletin & Review, 19(4), 555–574. 10.3758/s13423-012-0254-6

Shao, Z., Meyer, A. S., & Roelofs, A. (2013). Selective and nonselective inhibition of competitors in picture naming. Memory & Cognition, 41(8), 1200–1211. 10.3758/s13421-013-0332-7

Shao, Z., Roelofs, A., Martin, R. C., & Meyer, A. S. (2015). Selective inhibition and naming performance in semantic blocking, picture-word interference, and color–word Stroop tasks. *Journal of Experimental Psychology: Learning*, Memory, and Cognition, 41(6), 1806–1820. 10.1037/a0039363

Simon, J. R., & Rudell, A. P. (1967). Auditory S-R compatibility: The effect of an irrelevant cue on information processing. The Journal of Applied Psychology, 51(3), 300–304. 10.1037/h0020586

Stoffels, E. J., & van der Molen, M. W. (1988). Effects of visual and auditory noise on visual choice reaction time in a continuous-flow paradigm. Perception & Psychophysics, 44(1), 7–14. 10.3758/bf03207468

Strauss, E., Sherman, E., & Spreen, O. (2006). A compendium of neuropsychological tests: Administration, norms and commentary (3rd ed.). Oxford University Press.

Stroop, J. R. (1935). Studies of interference in serial verbal reactions. Journal of Experimental Psychology, 18(6), 643–662. 10.1037/h0054651

Tang, D., Chen, X., Li, H., & Lei, Y. (2022). Distributional analyses reveal the individual differences in congruency sequence effect. PLOS ONE, 17(8), e0272621. 10.1371/journal.pone.0272621

Vincent, S. B. (1912). The Function of the Vibrissae in the Behavior of the White Rat. University of Chicago.

Viviani, G., Di Pietro, I., Buchanan, E. M., Porru, A., Ambrosini, E., & Montefinese, M. (2025). Exploring semantic and executive flexibility interplay in task switching. Scientific Reports, 15(1), 31609. 10.1038/s41598-025-09639-y

Weissberger, G. H., Wierenga, C. E., Bondi, M. W., & Gollan, T. H. (2012). Partially overlapping mechanisms of language and task control in young and older bilinguals. Psychology and Aging, 27(4), 959–974. 10.1037/a0028281

Whitfield, K. E., Fillenbaum, G. G., Pieper, C., Albert, M. S., Berkman, L. F., Blazer, D. G., Rowe, J. W., & Seeman, T. (2000). The effect of race and health-related factors on naming and memory. The MacArthur Studies of Successful Aging. Journal of Aging and Health, 12(1), 69–89. 10.1177/089826430001200104

Wickham H, François R, Henry L, Müller K, Vaughan D (2023). dplyr: A Grammar of Data Manipulation. <10.32614/CRAN.package.dplyr>, R package version 1.1.4, <https://CRAN.R-project.org/package=dplyr>

Wylie, S. A., Ridderinkhof, K. R., Bashore, T. R., & van den Wildenberg, W. P. M. (2010). The effect of Parkinson’s disease on the dynamics of on-line and proactive cognitive control during action selection. Journal of Cognitive Neuroscience, 22(9), 2058–2073. 10.1162/jocn.2009.21326

Xiang, L., Zhang, B., Wang, B., Jiang, J., Zhang, F., & Hu, Z. (2016). The Effect of Aging on the Dynamics of Reactive and Proactive Cognitive Control of Response Interference. Frontiers in Psychology, 7. 10.3389/fpsyg.2016.01640

Yang, Q., & Pourtois, G. (2022). Reduced flexibility of cognitive control: Reactive, but not proactive control, underpins the congruency sequence effect. Psychological Research, 86(2), 474–484. 10.1007/s00426-021-01505-6

